# ZMYND11 Restrains KMT2A to Enable a Neuronal Developmental Program

**DOI:** 10.1101/2025.10.31.685692

**Authors:** Alexander W. Greben, Xiaoli S. Wu, Josephine E. Robb, Ian R. Vogel, Lisa Traunmüller, Sebastian Krüttner, Kara A. Fulton, Migue Van Louis Darcera, Eric C. Griffith, Michael E. Greenberg

## Abstract

Mutations in the chromatin reader and tumor suppressor *ZMYND11* are the cause of ZMYND11-related syndromic intellectual disability (ZRSID), a disorder characterized by symptoms such as language and motor delay, behavioral disruptions, and seizures. We find that neuronal deletion of *ZMYND11* in mice causes aberrant upregulation of non-neuronal gene programs, leading to reduced dendritic branching and spine density, as well as hyperactivity and abnormal motor behavior. We investigated the mechanism by which ZMYND11 regulates gene expression and discovered that ZMYND11 interacts with and inhibits the histone methyltransferase KMT2A (MLL1), a transcriptional co-activator which contributes to oncogenic and developmental gene programs. We find that a ZRSID-associated ZMYND11 point mutation abrogates ZMYND11 interaction with KMT2A, suggesting that this interaction is critical for the function of ZMYND11 in regulating brain development. Using a degron-tagged ZMYND11 mouse model to enable the rapid degradation of ZMYND11 in primary cortical neurons, we show that gene expression changes induced by ZMYND11 loss are attenuated by treatment with the KMT2A inhibitor revumenib, a drug which has recently been approved for the treatment of KMT2A-rearranged leukemia. Our findings shed light on the convergence of chromatin mechanisms regulating neuronal gene expression and raise the possibility that modulation of KMT2A activity may be a useful therapeutic avenue for ZRSID.

## Introduction

Mechanisms acting at the level of chromatin have emerged as a key site of dysfunction underlying neurodevelopmental disorders (NDD) and autism spectrum disorders (ASD). While cell fates are directed by the activity of lineage-determining transcription factors, the regulation of chromatin plays an essential role in promoting, restraining, or stabilizing transcriptional output^1,2^. Loss-of-function mutations in genes linked to chromatin regulation are increasingly implicated in a family of brain disorders termed chromatinopathies, with convergent features such as developmental delay, intellectual disability, and behavioral abnormalities^3,4^. In addition, mutations in many of the same chromatin regulators can lead to diseases such as cancer, indicating the importance of chromatin mechanisms for maintaining the precise orchestration of gene expression across tissues and developmental stages^5,6^.

The chromatin reader ZMYND11 was first linked to a neurodevelopmental disorder as a candidate gene for the 10p15.3 microdeletion syndrome, characterized by symptoms including intellectual disability, language and motor delay, behavioral abnormalities, and seizures^7–14^.

Subsequent genetic studies have confirmed that loss-of-function mutations in *ZMYND11* are the primary cause of developmental abnormalities in cases of 10p15.3 microdeletion, and have established a broader category of ZMYND11 gene disorders causing ZMYND11-related syndromic intellectual disability (ZRSID). Downregulation of *ZMYND11* expression is also linked to more aggressive cancer progression in a number of tissues, suggesting that ZMYND11 also functions as a tumor suppressor gene^15,16^. Consistent with ZMYND11’s function as a tumor suppressor, biochemical studies have revealed that the ZMYND11 protein binds and inhibits the viral transcriptional activators E1A and EBNA2 through a MYND-type zinc finger, which recognizes a degenerate PXLXP motif within these target proteins^17–21^. ZMYND11 has also been linked to inhibition of the splicing factor EFTUD2 and has been proposed to serve as a negative regulator of transcriptional elongation^22,23^. Supporting a role for ZMYND11 in neuronal development, a recent study found that ZMYND11 loss impairs the neuronal differentiation of neural stem cells, and that ZMYND11 regulates the expression of neuron-specific splice isoforms^24^.

Structural studies have shown that ZMYND11 is recruited to chromatin through a tripartite PHD-Bromo-PWWP domain that recognizes trimethylation at lysine 36 on the histone variant H3.3 (H3.3K36me3)^23^. The H3K36me3 mark is associated with active transcription, as its writer enzyme SETD2 binds to Ser2-phosphorylated RNA Polymerase II and co-transcriptionally modifies H3K36 with increasing density toward the 3’ end of expressed genes^25,26^. The H3.3 variant is itself enriched at sites of active transcription, and plays a unique role in post-mitotic cells such as neurons, making up the majority of all histone H3 as neurons mature^27,28^. Whether ZMYND11 bound to H3K36me3 across the transcribed regions of neuronal genes functions as an activator or repressor of transcription is not known, and more broadly, the mechanism by which ZMYND11 regulates gene expression in the brain has not previously been addressed.

Here we describe a key role for ZMYND11 in neuronal gene regulation and brain development. We find that mice with neuronal ablation of *Zmynd11* show stereotyped developmental abnormalities, including delayed growth and reduced dendritic branching and spine development in cortical pyramidal neurons. These mice also exhibit disruptions to behavior, including pronounced hyperactivity, repetitive behavior, and impaired motor coordination, all reminiscent of symptoms observed in humans bearing loss-of-function mutations in *ZMYND11*.

Investigating the mechanism by which ZMYND11 functions in the brain, we discovered that ZMYND11 interacts with and suppresses the activity of the chromatin regulator KMT2A (MLL1), a histone methyltransferase that promotes gene activation and plays essential roles in neuronal development. Like ZMYND11, the gene encoding KMT2A is mutated in a neurodevelopmental disorder, Wiedemann-Steiner Syndrome (WSS), and *KMT2A* is also frequently mutated in cancer^29–32^. The interaction of ZMYND11 with KMT2A is abrogated by a missense mutation within ZMYND11 (R600W) linked to ZRSID, suggesting that the interaction of ZMYND11 with KMT2A is critical for proper human brain development and that the disruption of this interaction underlies significant aspects of ZRSID. Finally, we established a degron mouse model in which endogenous ZMYND11 can be targeted for rapid degradation. Using this system, we demonstrate that loss of ZMYND11 induces changes in gene expression within hours which can be partially rescued by the KMT2A inhibitor revumenib, a drug recently approved for the treatment of KMT2A-rearranged leukemia^33,34^. These findings highlight the convergence of multiple molecular mechanisms which contribute to neurodevelopmental disorders, providing an advance toward a systemic understanding of chromatinopathies and new directions for the investigation of targeted therapeutics.

### ZMYND11 is essential for mouse viability and proper brain development

ZMYND11 is expressed across multiple tissues throughout development, but its expression is especially enriched in the nervous system **(Fig 1A)**. To investigate the function of ZMYND11 in the developing brain, we generated a *Zmynd11* conditional deletion mouse model **(Fig S1A)**. Using CRISPR-Cas9, we flanked exon 7 of *Zmynd11* with LoxP sites, which upon recombination introduces a frame-shift mutation and early stop codon within the *Zmynd11* gene.

**Figure 1.**
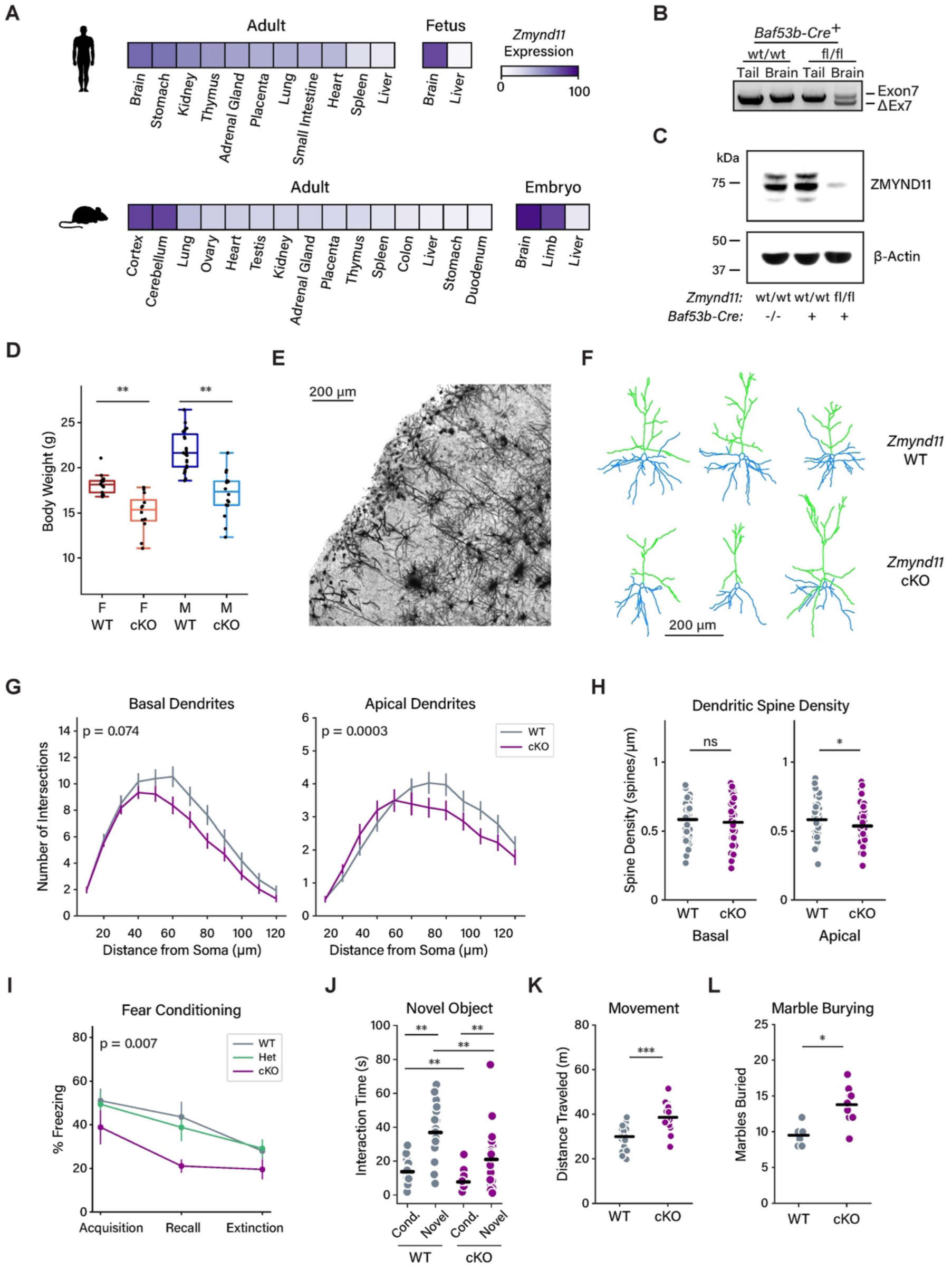
ZMYND11 regulates brain development and mouse behavior. (a) Heatmap showing relative levels of *Zmynd11* transcript in mouse (Mouse ENCODE transcriptome data) and human tissues (RNA sequencing of total RNA from 20 human tissues, UConn Health Center). (b) PCR genotyping showing Cre-mediated excision of *Zmynd11* exon 7 in *Zmynd11^fl/fl^* brain tissue with expression of *Baf53b-Cre*. (c) Western blot showing expression of ZMYND11 and β-Actin in cell lysates of cortical tissue. Zmynd11 protein is depleted in cortex from *Zmynd11^fl/fl^; Baf53b-Cre* (*Zmynd11-cKO*) mice. (d) Box plot showing the distribution of body mass of 6-week-old *Zmynd11^fl/fl^; Baf53b-Cre* (*Zmynd11-cKO*) mice and wild-type (*Zmynd11^wt/wt^; Baf53b-Cre*) littermates. ** = p < 0.001, p-values calculated using Wilcoxon rank-sum test, n = 13 WT Female, n = 11 cKO Female, n = 20 WT Male, and n = 12 cKO Male. (e) Representative brightfield image (minimum-intensity plot of confocal image stack) of Golgi-stained cortex from *Zmynd11-cKO* mouse at 10X magnification. Scale bar = 200 μm. Data are representative of 6 individual mice per condition. (f) Representative traces of individual Layer 2/3 cortical pyramidal neurons of 6-week-old *Zmynd11^fl/fl^; Baf53b-Cre* and *Zmynd11^wt/wt^; Baf53b-Cre* mice. Live tracing of brightfield images from Neurolucida software; scale bar = 200 μm. Data are representative of 6 cells per mouse, 6 individual mice per condition. (g) Sholl analysis of dendritic branch complexity in basal and apical dendrites of Layer 2/3 pyramidal neurons from 6-week-old *Zmynd11^fl/fl^; Baf53b-Cre* and *Zmynd11^wt/wt^; Baf53b-Cre* mice. Plot shows mean values ± SEM. p-values calculated using linear mixed-effects mode; n = 6 cells per mouse, 6 individual mice per condition. (h) Comparison of dendritic spine density in basal and apical dendrites of Layer 2/3 pyramidal neurons from 6-week-old *Zmynd11^fl/fl^;Baf53b-Cre* and *Zmynd11^wt/wt^; Baf53b-Cre* mice. * = p < 0.05, p-values calculated using Wilcoxon rank-sum test; n = 6 cells per mouse, 6 individual mice per condition. (i) Quantification of freezing behavior exhibited by *Zmynd11-cKO* mice and wild-type littermates (10-12 weeks old) upon contextual fear conditioning during acquisition (foot shock in novel context, final minute of 5-minute acquisition period), recall (repeat of context without additional foot shock, first 5 minutes in shock context cage), and extinction (after 25 minutes in shock context cage). p-values calculated using mixed model ANOVA, n = 7 individual mice (WT), n = 5 individual mice (Het), n = 8 individual mice (cKO). (j) Duration of mouse interactions with a novel and habituated object present in the same housing context, short-term recall. ** = p < 0.001, p-values calculated using Wilcoxon rank-sum test, n = 19 individual mice (WT), n = 21 individual mice (cKO). (k) Quantification of distance travelled by mice during the novel object recognition task in (j). *** = p < 0.0001, p-values calculated using Wilcoxon rank-sum test, n = 19 individual mice (WT), n = 21 individual mice (cKO). (l) Number of marbles buried by mice in open arena. * = p < 0.01, p-values calculated using Wilcoxon rank-sum test, n = 6 individual mice (WT), n = 9 individual mice (cKO).

As *ZMYND11* mutations linked to syndromic intellectual disability are deletion, truncation, or missense mutations predicted to be loss-of-function, the *Zmynd11*-*fl* mouse provides a useful model for understanding molecular and neurological abnormalities that occur in humans bearing mutations within *ZMYND11*. To model the human disorder that results from germline mutations, we first crossed *Zmynd11^fl/fl^* mice to the germline-expressing *E2a-Cre* **(Fig S1B)**. The resulting mice (*Zmynd11-KO*) exhibited germline recombination of exon 7 and lack detectible ZMYND11 protein **(Fig S1C,D)**. Levels of *Zmynd11* transcript were also reduced, suggesting nonsense-mediated decay of the recombined *Zmynd11-ΔExon7* transcript **(Fig S1E)**.

From crosses of *Zmynd11^+/-^* mice, we observed no homozygous knockout *Zmynd11^-/-^* pups across over 20 weaned litters **(Fig S1F)**. Examining late-gestation embryos from these crosses at E18.5, we observed *Zmynd11^-/-^* embryos at normal Mendelian ratios, suggesting homozygous loss of *Zmynd11* leads to early postnatal lethality. Conversely, *Zmynd11^+/-^* mice were viable and not readily distinguishable from wild-type littermates. RNA-Seq libraries from *Zmynd11^+/-^* cortical tissue also showed no abnormalities in gene expression compared to wild-type tissue aside from a 50% reduction in transcript levels of *Zmynd11* itself **(Fig S1G).** These findings suggest that, unlike humans, mice can readily tolerate the loss of a single copy of *Zmynd11*. The fact that homozygous loss of *Zmynd11* leads to early lethality in mice may also explain why no examples of homozygous mutation have been identified in humans.

Given the lethality of the homozygous deletion of *Zmynd11* and the lack of an evident phenotype upon heterozygous deletion, we took advantage of our conditional mouse model to generate a tissue-specific knockout of *Zmynd11*. We crossed *Zmynd11^fl/fl^* mice to the neuronal Cre driver *Baf53b-Cre*, which is expressed as neurons are specified during development, resulting in efficient recombination of the floxed exon and depletion of ZMYND11 protein within brain tissue **(Fig 1B,C, Fig S1H)**^35^. *Zmynd11^fl/fl^; Baf53b-Cre^+/^* (*Zmynd11-cKO)* mice are viable and survive into adulthood but show a pronounced reduction in their overall body size compared to littermates **(Fig 1D)**. To investigate changes in neuronal development, we used the Golgi method to stain brain tissue from *Zmynd11-cKO* mice and wild-type littermate controls at six weeks of age **(Fig 1E,F)**^36,37^. While cortical thickness was normal in *Zmynd11-*cKO mice despite their small overall size, morphological analysis of individual Layer 2/3 cortical neurons revealed a significant reduction in dendritic branch complexity in apical dendrites and a trend toward similar reduced branch complexity in basal dendrites **(Fig 1G, Fig S1I,J)**. We also observed a significant reduction in dendritic spine density in apical but not basal dendrites **(Fig 1H)**.

Together, these findings indicate that neuronal-specific deletion of *Zmynd11* leads to morphological abnormalities in cortical neurons in the mouse brain, which may contribute to observed developmental abnormalities such as delayed growth.

We next performed behavioral testing to assess whether *Zmynd11-cKO* mice can serve as a model for studying symptoms of ZRSID. In a contextual fear conditioning paradigm, when compared to wild-type mice *Zmynd11-cKO* mice showed reduced freezing behavior in response to foot shock, which persisted during recall **(Fig 1I, Fig S2A)**. In a test of novel object recognition, *Zmynd11-cKO* mice displayed reduced interaction with both familiar and novel objects, but a similar relative preference for novelty **(Fig 1J, Fig S2B,C)**. Taken together, this does not indicate a clear deficit in memory, but may reflect a level of heightened activity that led *Zmynd11-*cKO mice to freeze less and to spend less time investigating test objects compared to wild-type littermates. Quantification of total distance traveled during novel object recognition testing confirmed a heightened level of movement in *Zmynd11-cKO* mice. In a marble-burying assay, a measure of repetitive behaviors such as those often found in ASD, *Zmynd11-cKO* mice buried significantly more marbles than wild-type littermates **(Fig 1K,L)**.

These findings of hyperactivity and repetitive behaviors in *Zmynd11-cKO* mice are consistent with observations in humans with ZRSID, which is associated with hyperactivity and attention-deficit hyperactivity disorder (ADHD)^7,11–14^. We also observed deficits in motor behavior in *Zmynd11-cKO* mice, which fail to maintain balance on a rotating rod compared to wild-type littermates **(Fig S2D)**. By contrast, wild-type and *Zmynd11-cKO* mice showed similar levels of exploration of open areas in an elevated plus maze, indicating that *Zmynd11-cKO* mice do not display an obvious anxiety phenotype **(Fig S2E)**. Taken together, the behavioral disruptions we observed in *Zmynd11-cKO* mice are reminiscent of the clinical features observed in individuals with ZRSID, indicating that *Zmynd11-cKO* mice can serve as a useful model for understanding the pathogenesis of ZRSID and for the evaluation of therapies to treat this disorder.

### ZMYND11 regulates neuronal gene expression

To investigate the effect of ZMYND11 loss on neuronal gene expression, we used an adeno-associated virus (AAV) to deliver Cre or a nonfunctional truncated form of Cre (ΔCre) to opposite cortical hemispheres of *Zmynd11^fl/fl^* mice **(Fig 2A)**. RNA libraries prepared from *Zmynd11^fl/fl^; AAV-Cre* cortex showed a reduction in *Zmynd11* transcripts relative to the *ΔCre* control, and RNA-Seq analysis of gene expression revealed a widespread dysregulation of neuronal gene transcription when ZMYND11 function is disrupted **(Fig 2B)**. Upon depletion of ZMYND11 more than a thousand genes are significantly up- and down-regulated, with up- regulated genes being more numerous and showing a greater magnitude change compared to down-regulated genes. Genes down-regulated in *Zmynd11-cKO* were enriched for roles in neuronal development and synaptic transmission, whereas up-regulated genes were enriched for roles in non-neuronal cellular functions **(Fig 2C)**. Specifically, upregulated genes were enriched for roles in Wnt signaling, MAPK signaling, and pathways linked to cancer **(Fig S3A,B)**. These findings suggest that ZMYND11 enables a pro-neuronal developmental gene program, whether by promoting the expression of neuronal genes or inhibiting the aberrant expression of non-neuronal genes.

**Figure 2.**
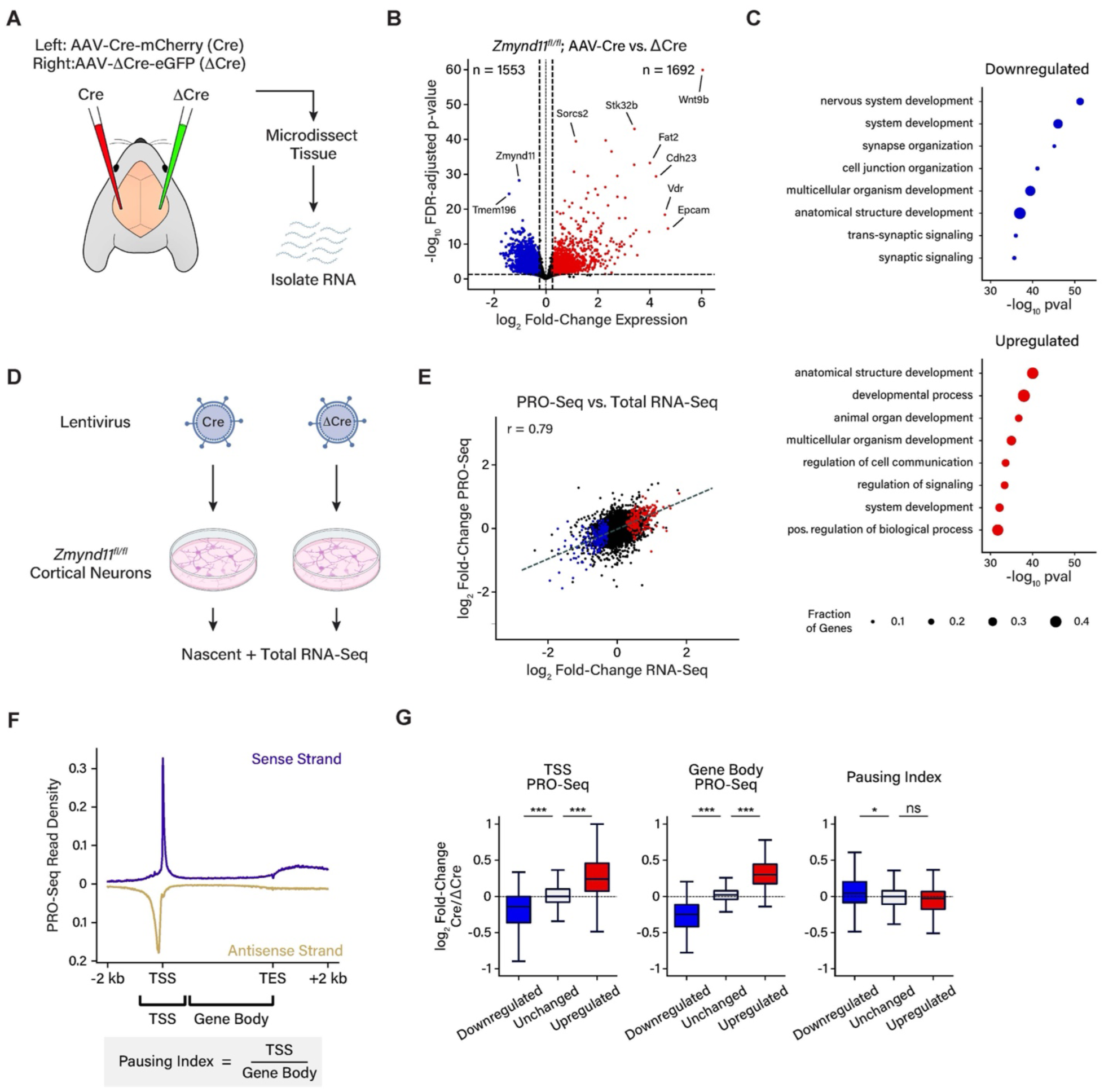
Regulation of neuronal gene expression by ZMYND11. (a) Schematic showing stereotactic injection of AAV expressing either Cre recombinase or non-functional ΔCre into opposite hemispheres of primary visual cortex (V1) of 4-week-old *Zmynd11^fl/fl^* mice. (b) Volcano plot of gene expression changes in *Zmynd11^fl/fl^* AAV-Cre vs. AAV-ΔCre cortex. Plot shows DESeq2 log_2_ fold-change RNA expression vs. -log_10_ p-value with Benjamini-Hochberg false discovery rate (FDR) adjustment. Dashed lines show cutoff for log_2_ fold-change (± 0.25) and FDR-adjusted p-value (0.1). Tissue from primary visual cortex (n = 8 individual mice). (c) Gene ontology analysis of DESeq2 significantly up- and down-regulated genes in *Zmynd11^fl/fl^* AAV-Cre vs. AAV-ΔCre cortex, GO:BP (Biological Process) gene set, analysis by gProfiler2. (d) Schematic showing infection of DIV7 primary cortical neurons from *Zmynd11^fl/fl^* embryos with lentivirus expressing either Cre or ΔCre. Samples were collected for total RNA and nascent RNA (PRO-Seq) processing. (e) Scatter plot showing changes in total and nascent RNA expression in *Zmynd11^fl/fl^* Lenti-Cre vs. Lenti-ΔCre DIV 7 cortical cultures. Plot shows log_2_ fold-change total RNA-seq vs. log_2_ fold-change PRO-Seq; dotted line shows linear fit with Pearson correlation coefficient labeled. n = 3 independent neuron dissections (RNA-Seq), and n = 4 independent neuronal dissections (PRO-Seq). (f) Aggregate plots showing the average distribution of PRO-Seq signal across genes in wild-type primary cortical neurons. Gene body and transcription start site (TSS) regions highlighted for computation of Pausing Index. (g) Box plot of PRO-seq counts at the transcription start site (left) and gene body (middle) of significant upregulated, downregulated, and unchanged genes by DESeq2 in *Zmynd11^fl/fl^* Lenti-Cre vs. Lenti-ΔCre. (Right) Box plot showing pausing index (ratio of PRO-Seq counts at TSS/GB) at up-, down-, and unchanged genes. *** = p < 1E-30, * = p < 1E-05, p-values calculated by Wilcoxon rank-sum test (n = 4 independent neuronal dissections).

Prior work in non-neuronal cells has highlighted the function of ZMYND11 in regulating mRNA splicing, showing that ZMYND11 promotes intron retention by inhibiting EFTUD2^22^. We similarly found that splicing is dysregulated in *Zmynd11-cKO* cortex, with a bias toward retention of alternate exons as well as introns, indicating that ZMYND11 regulates mRNA splicing in the brain **(Fig S4A-C)**. However, in the absence of ZMYND11, many more genes display a significant change in their overall expression than a change in splicing. Indeed, genes that displayed an alteration in splicing in *Zmynd11-cKO* mice were no more likely to show significant changes in their expression than genes whose splicing was unaffected, suggesting that ZMYND11 may regulate gene expression and RNA splicing by independent mechanisms **(Fig S4D)**.

ZMYND11 has also been proposed to be an inhibitor of transcriptional pause release in a cancer cell line, as shRNA-mediated knockdown of *ZMYND11* in U2OS cells leads to an increase in the occupancy of RNA Polymerase II at gene bodies relative to the promoter of upregulated genes^23^. To test whether this is a generalizable mechanism of gene repression by ZMYND11, we disrupted ZMYND11 on a shorter timescale to reduce the impact of secondary effects due to widespread misregulation of gene expression **(Fig 2D)**. Towards this end we cultured primary cortical neurons from *Zmynd11^fl/fl^* mice and infected them with lentivirus expressing Cre or ΔCre for 5 days. At this shorter time point, we found by RNA-Seq that when ZMYND11 function is disrupted, the expression of several hundred genes is significantly up- or down-regulated **(Fig S3C)**. To observe nascent transcription at high resolution, we used Precision Run-On Sequencing (PRO-Seq) to map the position of actively elongating RNA Pol II across the genome^38^. We found that genes which show significant increase or decrease in total RNA levels in *Zmynd11^fl/fl^;Lenti-Cre* show a corresponding increase or decrease in the level of their nascent RNA transcripts **(Fig 2E**), indicating that the observed changes in transcript abundance are indeed due to changes in the level of transcription, rather than transcript processing or stability. The relative enrichment of PRO-Seq signal can also be compared between promoters and gene bodies to assess the relative rate of transcriptional pausing compared to productive elongation, a measure known as the pausing index **(Fig 2F)**. We observed that changes in PRO-Seq signal at promoters and gene bodies were largely correlated, likely reflecting a change in transcriptional initiation rather than pause release. A subset of the down-regulated genes showed a mild increase in the pausing index, while there was no significant change in the pausing index of up-regulated genes **(Fig 2G, S3D)**. This finding suggests that regulation of transcriptional pause release is not a general mechanism by which ZMYND11 regulates transcription, and if ZMYND11 is indeed a regulator of transcriptional elongation, this function may differ across cell types or developmental contexts. Taken together, our results suggests a role for ZMYND11 in transcriptional regulation outside of splicing or pausing. In particular, ZMYND11 appears to be essential for the maintenance of proper neuronal gene expression, with its loss leading to the upregulation of gene programs associated with Wnt signaling, non-neuronal development, and oncogenesis.

### ZMYND11 binds active chromatin across the genome

To investigate the mechanism by which ZMYND11 regulates neuronal gene expression, we next examined the distribution of ZMYND11 across the neuronal genome using CUT&RUN to map ZMYND11 occupancy in cortical tissue from wild-type and *Zmynd11-cKO* mice. In wild-type cortex, we observed ZMYND11 binding across thousands of expressed genes, with an enrichment at both the 5’ and 3’ ends of genes **(Fig 3A)**. The binding of ZMYND11 across gene bodies is strongly correlated with the presence of H3K36me3 across these regions, and ZMYND11, like H3K36me3, is enriched across gene bodies in proportion with the level of a gene’s expression. However, we unexpectedly found that ZMYND11 also binds strongly to the transcription start site of actively expressed genes, a finding that is unlikely to be explained by interactions with H3K36me3, as this mark is strongly depleted at transcription start sites.

**Figure 3.**
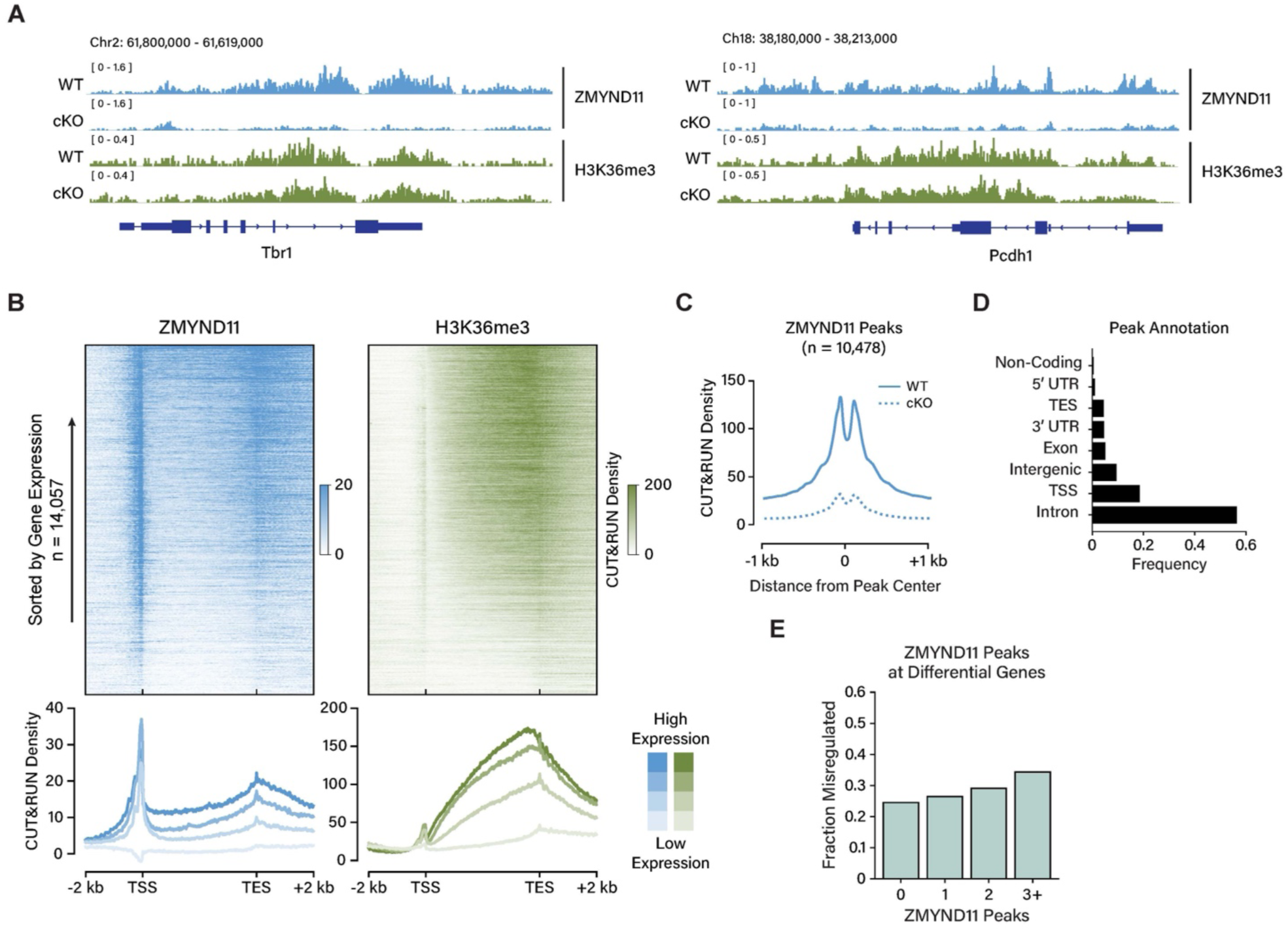
ZMYND11 binds active chromatin across the genome. (a) Representative tracks from IGV (Integrated Genomics Viewer) showing CUT&RUN signal for ZMYND11 and H3K36me3 in wild-type (*Zmynd11^wt/wt^; Baf53b-Cre*) and *Zmynd11-cKO* (*Zmynd11^fl/fl^; Baf53b-Cre*) cortex at two example loci. (b) (Above) Heatmaps of CUT&RUN signal from cortical tissue (scaled read coverage per bp) across all genes for ZMYND11 and H3K36me3. (Below) Aggregate plots showing the average distribution of CUT&RUN signal for ZMYND11 and H3K36me3 across genes separated into quartiles by level of expression. ZMYND11 CUT&RUN is normalized by subtraction of signal present in *Zmynd11-cKO*; n = 2 independent replicates per condition. (c) Aggregate plots showing average distribution of ZMYND11 CUT&RUN signal centered around significant ZMYND11 peaks in *Zmynd11^fl/fl^;Baf53b-Cre* (cKO) and *Zmynd11^wt/wt^; Baf53b-Cre* (WT) brain tissue (MACS2 no-control p-value < 1e-05, ZMYND11 CUT&RUN WT/cKO > 2). (d) Annotation of the genomic localization of ZMYND11 peaks by Homer annotatePeaks (MACS no-control p-value < 1e-05, ZMYND11 CUT&RUN WT/cKO > 2). (e) Proportion of genes with significant differential expression in *Zmynd11^fl/fl^* AAV-Cre vs. AAV-ΔCre, grouped by the number of ZMYND11 CUT&RUN peaks per gene.

Importantly, in *Zmynd11-cKO* mice there is strong depletion of ZMYND11 CUT&RUN signal at both gene body and transcription start sites, demonstrating the specificity of ZMYND11 CUT&RUN signal **(Fig 3B)**. Collectively, we identified 10,476 ZMYND11 peaks showing greater than two-fold enrichment in wild-type over cKO, 70% of peaks lying within gene bodies and 20% at transcription start sites **(Fig 3C,D)**.

Notably, the number of significant ZMYND11 peaks occurring within a gene is predictive of the likelihood of that gene being misregulated in *Zmynd11^fl/fl^ AAV-Cre-*treated cortex, but the number of peaks is not predictive of whether the gene will be up- or down-regulated **(Fig 3E, S5A)**. Thus, while ZMYND11 binds broadly across the genome at thousands of expressed genes, the binding of ZMYND11 at a gene is not sufficient to determine whether the gene will be up- or down-regulated upon loss of ZMYND11. This finding could indicate that ZMYND11 can positively or negatively regulate gene expression, or it could instead reflect primary and secondary effects of a broadly dysregulated transcriptional state upon *Zmynd11* deletion.

The level of expression of a given gene was the most predictive genomic feature for the degree of ZMYND11 binding at the gene, whether at the start site, end site, or gene body, with high levels of ZMYND11 binding correlating with a high level of gene expression **(Fig S5B)**.

ZMYND11 binding at gene bodies is highly correlated with the presence of the histone mark H3K36me3, but also with other histone modifications found at active genes, such as H3K27 acetylation, and mono- and tri-methylation of H3K4. By contrast, ZMYND11 peaks at transcription start sites or within intergenic regions lack H3K36me3 but showed other signatures of active transcription such as H3K27ac and H3K4me3 **(Fig S5C)**. These dichotomous binding sites suggest a link between ZMYND11 and active chromatin that is broader than the previously identified function of ZMYND11 as a reader of H3K36me3. The absence of the H3K36me3 mark in promoter regions makes this mark unlikely to be the determinant of ZMYND11 localization at these sites, indicating other factors likely contribute to ZMYND11 binding at promoters.

CUT&RUN reads for ZMYND11 were largely nucleosomal (>150 bp), supporting a mode of genomic binding in which ZMYND11 largely interacts with chromatin rather than nucleosome-depleted DNA **(Fig S5D)**. Further supporting this interpretation, no consistent DNA motif can be found within ZMYND11 peaks, indicating that ZMYND11 binding is not directed by recognition of a defined sequence motif within DNA **(Fig S5E)**. Rather, ZMYND11 peaks at transcription start sites and within gene bodies are enriched for the binding sites of a variety of common transcription factors, such as ETS and NFY at promoters, and MEF2 and EGR within gene bodies. This mode of binding is consistent with the known function of ZMYND11 as a reader of active chromatin, with H3K36me3 and additional chromatin features possibly contributing to ZMYND11 localization.

### ZMYND11 interacts with KMT2A in the brain

Having identified a clear role for ZMYND11 in the regulation of neuronal transcription, we sought to investigate the mechanism by which ZMYND11 binding regulates expression of target genes. ZMYND11 was first identified as a repressor of the adenovirus transcriptional activator E1A, and transcriptional reporter assays in cell lines have shown that ZMYND11 functions as a repressor of viral as well as cellular transcription factors, including EBNA2 and c-Myb^17,19,39^.

However, while we found that ZMYND11 regulates neuronal transcription, the molecular mechanisms underlying this function are not explained by previously characterized interactions of ZMYND11 with other proteins.

Because ZMYND11 lacks domains with predicted enzymatic activity, we considered the possibility that it represses or activates target genes through the recruitment of cofactors, or by direct inhibition through binding target proteins. We therefore pursued an immunoprecipitation- mass spectrometry (IP-MS) approach to purify ZMYND11 from brain tissue and identify its interacting partners. To enable efficient immunoprecipitation of endogenous ZMYND11 from neuronal chromatin without disrupting potential interactors, we generated a knock-in mouse model in which a FLAG-HA tag is fused to the N-terminus of ZMYND11 **(Fig 4A, Fig S6A)**.

**Figure 4.**
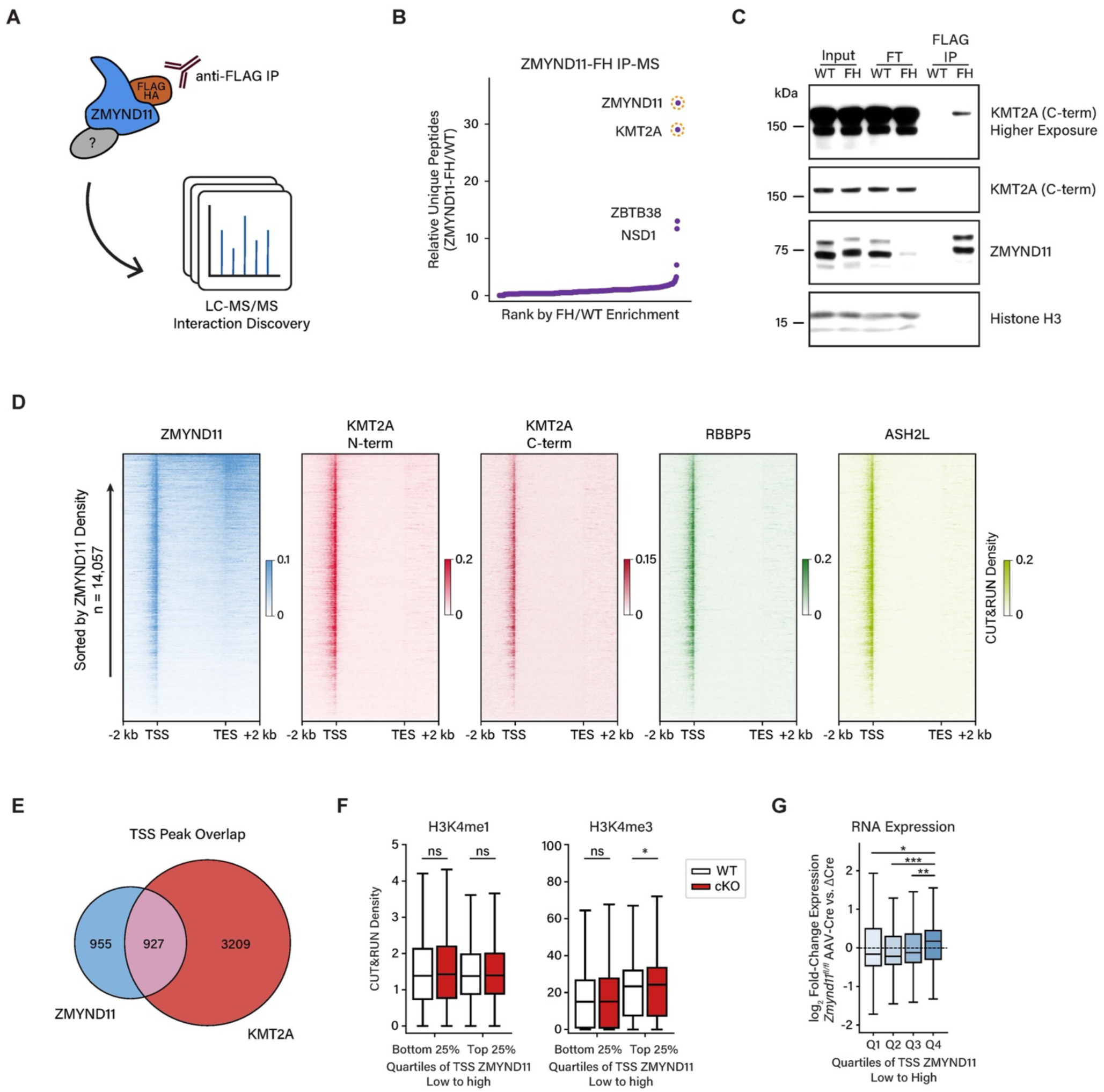
ZMYND11 interacts with KMT2A in the brain. (a) Schematic showing experimental approach of immunoprecipitation followed by protein mass spectrometry for the discovery of proteins interacting with endogenously FLAG-HA-tagged ZMYND11. (b) Scatter plot showing proteins identified by mass spectrometry in eluates from anti-FLAG immunoprecipitation of ZMYND11-FH from whole brain tissue, ranked by enrichment in ZMYND11-FH tissue over wild-type; n = 3 independent replicates per condition. (c) Representative western blot showing co-immunoprecipitation (co-IP) of ZMYND11 and KMT2A from nuclear lysates of brain tissue from *Zmynd11^FH/FH^* (FH) and wild-type littermates (WT). FT: Flow-through, IP: anti-FLAG immunoprecipitation. Data are representative of 3 independent experiments. (d) Heatmap of CUT&RUN signal (scaled read coverage per bp) across genes ranked by ZMYND11 occupancy for KMT2A (N- and C-terminal fragments), RBBP5, ASH2L, and ZMYND11 in 6-week-old mouse cortex; n = 3 individual mice per condition. (e) Venn diagram showing overlap of TSS-localized ZMYND11 peaks and MLL1 consensus peaks. (f) Box plots showing H3K4 mono- (H3K4me1) and tri-methylation (H3K4me3) levels at top and bottom quartiles of ZMYND11-bound transcription start sites, showing the difference in CUT&RUN density between wild-type and *Zmynd11-cKO* in 6-week-old mouse cortex. * = p < 0.01, p-values calculated using Wilcoxon rank-sum test; n = 3 independent replicates per condition. (g) Box plots showing differential gene expression by DESeq2 analysis of RNA-seq in *Zmynd11^fl/fl^* AAV-Cre vs. AAV-ΔCre, with genes separated into quartiles based on enrichment of WT/cKO ZMYND11 CUT&RUN signal at TSS (low to high). * = p < 0.05, ** = p < 0.01, *** = p < 0.00001, p-values calculated using Wilcoxon rank-sum test; n = 3 independent replicates per condition.

Homozygous *Zmynd11-FH* mice were viable and healthy, and tagged ZMYND11 protein mapped by anti-FLAG CUT&RUN showed highly similar genomic localization to wild-type ZMYND11 **(Fig S6B)**. ZMYND11 peaks that were initially defined by sensitivity to *Zmynd11-cKO* showed corresponding FLAG CUT&RUN signal in *Zmynd11^FH^* cortical tissue which is depleted in wild-type mice, confirming the genomic binding of ZMYND11-FH protein and the specificity of CUT&RUN signal for endogenous and tagged ZMYND11 **(Fig S6C,D)**. Together, these studies show that epitope tagging of ZMYND11 does not disrupt ZMYND11 expression or genomic localization, making the tagged protein a useful tool for identifying endogenous ZMYND11-interacting proteins.

Using antibodies recognizing the FLAG epitope tag, we immunoprecipitated FLAG-HA-tagged ZMYND11 from nuclear lysates of *Zmynd11-FH* brain tissue and subjected eluates to mass spectrometry to identify proteins co-immunoprecipitating with ZMYND11. We identified several protein-protein interactions with endogenous ZMYND11, most prominently the histone methyltransferase KMT2A (also known as MLL1), the core subunit of a transcriptional regulatory complex linked to H3K4 trimethylation and gene activation **(Fig 4B, S6E,F)**. KMT2A has been extensively studied for its prominent roles in cancer as well as in brain development, where mutations in the gene *KMT2A* cause the neurodevelopmental disorder Wiedemann-Steiner Syndrome^29,32^. We also identified the H3K36 dimethyltransferase NSD1 and the methyl-DNA-binding transcription factor ZBTB38 as ZMYND11-interacting proteins^40^. NSD1 is a chromatin-modifying factor linked to transcriptional regulation and a neurodevelopmental disorder, as mutations in the *NSD1* gene cause Sotos Syndrome^41,42^. Due to the high level of enrichment of KMT2A in ZMYND11 immunoprecipitates and its well-characterized links to transcriptional regulation and brain development, we subsequently focused on KMT2A as a candidate ZMYND11 interactor of interest.

We first validated co-immunoprecipitation between ZMYND11 and KMT2A in *Zmynd11-FH* brain tissue by western blot **(Fig 4C)**. We further characterized this interaction by expressing FLAG-HA-tagged ZMYND11 in HEK293T cells, confirming that ZMYND11 co-immunoprecipitates with KMT2A as well as with additional known MLL1 complex members such as RBBP5, WDR5, and MENIN **(Fig S7A)**. Notably, interactions between ZMYND11 and MLL1 complex members were reduced by treatment with ethidium bromide, suggesting that the ZMYND11-KMT2A interaction is partially dependent on the presence of DNA, potentially reflecting a preference of ZMYND11 for interacting with KMT2A in the context of chromatin.

ZMYND11 showed much less interaction with KMT2B, a homolog of KMT2A, and did not show significant interaction with TBP or CREB, suggesting that ZMYND11 is not simply binding non-specifically to factors at active promoters.

To determine whether ZMYND11 and KMT2A co-localize on chromatin, we mapped the binding of MLL1 core complex members KMT2A, RBBP5, and ASH2L to chromatin in mouse cortex by CUT&RUN **(Fig 4D)**. We observed a high level of co-occupancy of MLL1 complex members and identified 7440 consensus MLL1 peaks **(Fig S7B)**. As expected for MLL1, these peaks preferentially localize to the transcription start sites of active genes **(Fig S7C)**. Notably, we observed a high level of ZMYND11 co-occupancy at MLL1 binding sites, with half of ZMYND11 peaks found within TSS regions directly overlapping a KMT2A peak **(Fig 4E)**. This striking degree of overlap between ZMYND11 and the KMT2A/MLL1 complex suggests that ZMYND11 localized at transcription start sites may frequently be interacting with KMT2A.

We went on to examine histone methylation within the top quartile of ZMYND11-bound transcription start sites and found no change in H3K4me1 in *Zmynd11-cKO* cortical tissue compared to wild-type mice. However, in *Zmynd11-cKO*, we found a significant increase in H3K4me3 levels at sites highly bound by ZMYND11, but not at sites lowly bound by ZMYND11 **(Fig 4F)**. In addition, we found that genes exhibiting strong ZMYND11 binding at their transcription start sites show increased expression upon *Zmynd11* disruption. By contrast, we observed no correlation between the enrichment of ZMYND11 at gene bodies or transcription end sites and gene up-regulation in *Zmynd11-cKO* **(Fig 4G, S7D)**. Together, these results suggest that promoter-bound ZMYND11 represses transcription, and raise the possibility that ZMYND11 binding inhibits KMT2A and suppresses KMT2A-dependent transcriptional activation.

### Disease-associated *ZMYND11* mutations disrupt interactions with KMT2A

To determine if the interaction of ZMYND11 with KMT2A affects ZMYND11 function, we next used disease-associated mutations as a lens through which to probe ZMYND11 molecular function. We reasoned that if a mutation of *ZMYND11* that leads to ZRSID also disrupts the interaction between ZMYND11 and KMT2A, this would indicate the functional importance of the interaction. The MYND domain of ZMYND11 is a hotspot of mutations within the *ZMYND11* gene that are linked to ZRSID, with the most observed missense mutation being an arginine-to-tryptophan substitution at amino acid position 600 (R600W). This mutation occurs at the C-terminal end of the MYND domain at a residue found to be essential for interaction with the adenovirus E1A protein **(Fig 5A)**^43^. In addition to the R600W mutation, partial deletions of the MYND domain are also associated with ZRSID, suggesting that key functions of ZMYND11 are dependent on this domain **(Fig S8A)**.

**Figure 5.**
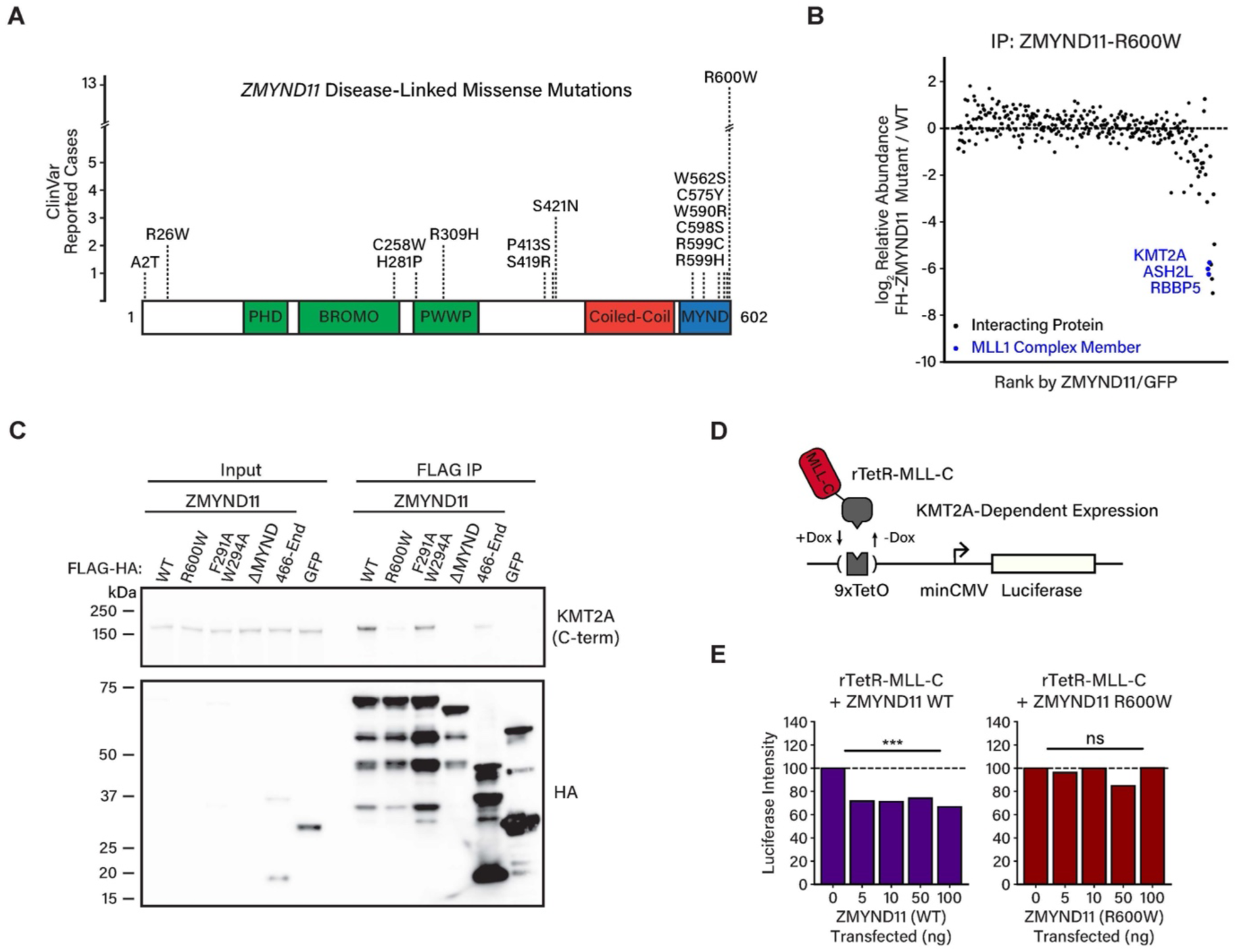
Disease-associated *ZMYND11* mutations disrupt interactions with KMT2A. (a) Schematic showing missense mutations found in cases of *ZMYND11*-linked neurodevelopmental disorder in human patients, from ClinVar reports evaluated ‘Pathogenic’ and ‘Likely Pathogenic’. (b) Scatter plot showing the relative enrichment of co-immunoprecipitating proteins with ZMYND11-R600W compared to wild-type ZMYND11, ordered by relative enrichment in wild-type vs. FLAG-tagged GFP control. KMT2A (MLL1) complex members are indicated in blue; n = 3 independent transfections per condition. (c) Representative western blot showing co-immunoprecipitation of FLAG-HA-tagged ZMYND11 and endogenous KMT2A. ZMYND11 constructs with a dual FLAG-HA tag are expressed in HEK293T cells, and nuclear lysates subjected to immunoprecipitation with anti-FLAG antibodies. Anti-HA western blot shows purification of bait protein, while anti-KMT2A (C-term) western blot shows co-IP of interacting protein. Data are representative of three independent experiments. (d) Schematic showing implementation of luciferase reporter assay for KMT2A-dependent gene activation. (e) Bar plots showing relative luminance intensity, ratio of firefly luciferase (expressed by test plasmid) to renilla luciferase (expressed by co-transfected control plasmid) in HEK293T cells expressing *9xTetO-minCMV-Luciferase* reporter and *rTetR-MLL-C* with or without co-expression of wild-type ZMYND11 (left) or ZMYND11-R600W (right). *** = p < 0.001, p-values by two-sided independent samples t-test; n = 2 independent transfections per condition, two independent measurements per sample.

To assess whether mutations within the MYND domain affect the interaction of ZMYND11 with KMT2A or other proteins, we expressed various FLAG-tagged ZMYND11 constructs in HEK293T cells, then immunoprecipitated ZMYND11 with an anti-FLAG antibody and identified ZMYND11-interacting proteins by quantitative tandem mass tag mass spectrometry (TMT-MS) ^44,45^. In this analysis, we compared wild-type ZMYND11 to the R600W mutant, as well as a truncated ZMYND11 mutant lacking the MYND domain (ΔMYND) **(Fig S8B,C)**. We found that ZMYND11 expressed in 293T cells co-immunoprecipitated with proteins such as KMT2A and NSD1 that we also identified in the mouse brain, in addition to a wider range of transcriptional regulators and chromatin-associated proteins that may reflect lower-affinity interactions observed due to ZMYND11 overexpression **(Fig S8D,E)**. Deletion of the MYND domain (ZMYND11-ΔMYND) led to a drastic reduction in the majority of proteins interacting with ZMYND11, whereas the ZMYND11-R600W mutant showed a more selective loss of the top ZMYND11 interactors, including KMT2A and NSD1 as well as MLL1 complex members ASH2L and RBBP5 **(Fig 5B, Fig S8F)**.

We next characterized in more detail the regions of ZMYND11 that are necessary and sufficient for interaction with KMT2A by immunoblotting ZMYND11 immunoprecipitation eluates from 293T cells expressing wild-type or mutant ZMYND11. We confirmed that the ZMYND11-R600W mutation, as well as the ZMYND11-ΔMYND truncation, significantly decreased the interaction of ZMYND11 with KMT2A **(Fig 5C)**. By contrast, a minimal C-terminal region of ZMYND11, corresponding to the MYND domain and flanking sequences, effectively co-immunoprecipitated with KMT2A, although to a lesser degree than full-length ZMYND11.

Notably, we found that ZMYND11 mutations (F291A+W294A) which lie within the ZMYND11 PWWP domain and prevent binding of ZMYND11 to H3K36me3 did not affect ZMYND11 binding to KMT2A, suggesting that ZMYND11 interactions with KMT2A occur independently of binding to H3K36me3. Together, these findings demonstrate the necessity and partial sufficiency of the ZMYND11 MYND domain for the interaction with KMT2A and confirm that the disease-associated R600W mutation strongly disrupts this interaction.

### ZMYND11 represses KMT2A-dependent transcription

We next asked if the binding of ZMYND11 to KMT2A promotes transcriptional activation, or whether it inhibits the ability of KMT2A to activate transcription. Toward this end, we used a K562 reporter cell line in which a Citrine fluorescent reporter gene is expressed under the control of a promoter containing multiple TetO-binding sites (*9xTetO-pEF-Citrine*), which are bound by the rTetR protein domain in the presence of doxycycline **(Fig S9A)**^46^. We first introduced into these cells an rTetR fusion of the ZMYND11 MYND domain (rTetR-MYND) and examined the effect on reporter gene expression **(Fig S9B)**. Recruitment of rTetR-MYND in *9xTetO-pEF-Citrine* cells led to the repression of the reporter gene, as measured by a reduction in the proportion of reporter cells expressing Citrine **(Fig S9C)**. In contrast, an rTetR-MYND fusion bearing the R600W mutation, which fails to bind KMT2A, was significantly less effective at repressing *9xTetO-pEF-Citrine* reporter gene transcription **(Fig S9D)**, suggesting that R600W-sensitive interactions, such as the interaction between ZMYND11 and KMT2A, contribute to the repressive function of the ZMYND11 MYND domain.

We next used a reporter cell line employing a weak TetO-containing promoter that does not yield measurable basal levels of Citrine expression (*9xTetO-minCMV-Citrine)*, testing the potential ability of rTetR-MYND to potentiate reporter gene transcription. Recruitment of rTetR-MYND in *9xTetO-minCMV-Citrine* cells did not stimulate Citrine expression, whereas recruitment of the C-terminal fragment of KMT2A (rTetR-MLL-C) was sufficient to activate Citrine expression within a population of cells **(Fig S9E)**. These findings show that despite the capacity to bind endogenous KMT2A, the ZMYND11 MYND domain serves to repress, but not to activate reporter gene transcription, further evidence that ZMYND11 functions as a repressor of KMT2A.

Finally, to test the ability of full-length ZMYND11 to inhibit KMT2A-dependent transcription, we generated a reporter construct expressing a firefly luciferase gene driven by a weak promoter containing TetO-binding sites (*9xTetO-minCMV-Luciferase)* and transfected it into the HEK293T cell line. While these cells did not express measurable basal levels of luciferase, co-expression of an rTetR fusion of the C-terminal region of KMT2A (rTetR-MLL-C) with the luciferase reporter construct led to induction of luciferase expression in the presence of doxycycline, rendering this a useful reporter assay for KMT2A-dependent gene activation **(Fig 5D)**. Co-expression of full-length ZMYND11 in these cells significantly repressed luciferase expression driven by rTetR-MLL-C, whereas expression of ZMYND11-R600W had no effect on reporter expression **(Fig 5E)**. Thus, ZMYND11 binding to KMT2A through its MYND domain inhibits the ability of KMT2A to activate transcription, providing a mechanistic basis for the repression of gene expression by ZMYND11 and a causal link between pathogenic variants of ZMYND11 and defects in molecular function.

### Rapid Depletion of ZMYND11 and Rescue by KMT2A Inhibition

We next sought to investigate ZMYND11-dependent inhibition of KMT2A in the context of endogenous gene expression in neurons. However, the profound cascading dysregulation of neuronal gene programs accompanying genetic ablation of ZMYND11 complicates the study of the direct effects of ZMYND11 loss on target gene expression. To enable selective disruption of ZMYND11 with improved temporal resolution, we generated a knock-in mouse model in which the FKBP12^F36V^ inducible degron tag was fused to the N-terminus of ZMYND11 (*Zmynd11-dTAG*), enabling targeted degradation of ZMYND11 protein in the presence of heterobifunctional dTAG small molecules such as dTAG-13 and dTAG^v^-1 **(Fig 6A, Fig S10A)**^47,48^. Primary cortical neurons cultured from *Zmynd11-dTAG* animals showed rapid degradation of ZMYND11 protein after dTAG-13 treatment, with protein levels reduced within 15 minutes and nearly undetectable by 60 minutes **(Fig 6B)**. In the context of chromatin-bound ZMYND11, CUT&TAG analysis after three hours of dTAG-13 treatment revealed a dramatic reduction of ZMYND11 signal at over a thousand genes and hundreds of transcription start sites **(Fig 6C, Fig S10B,C)**.

**Figure 6.**
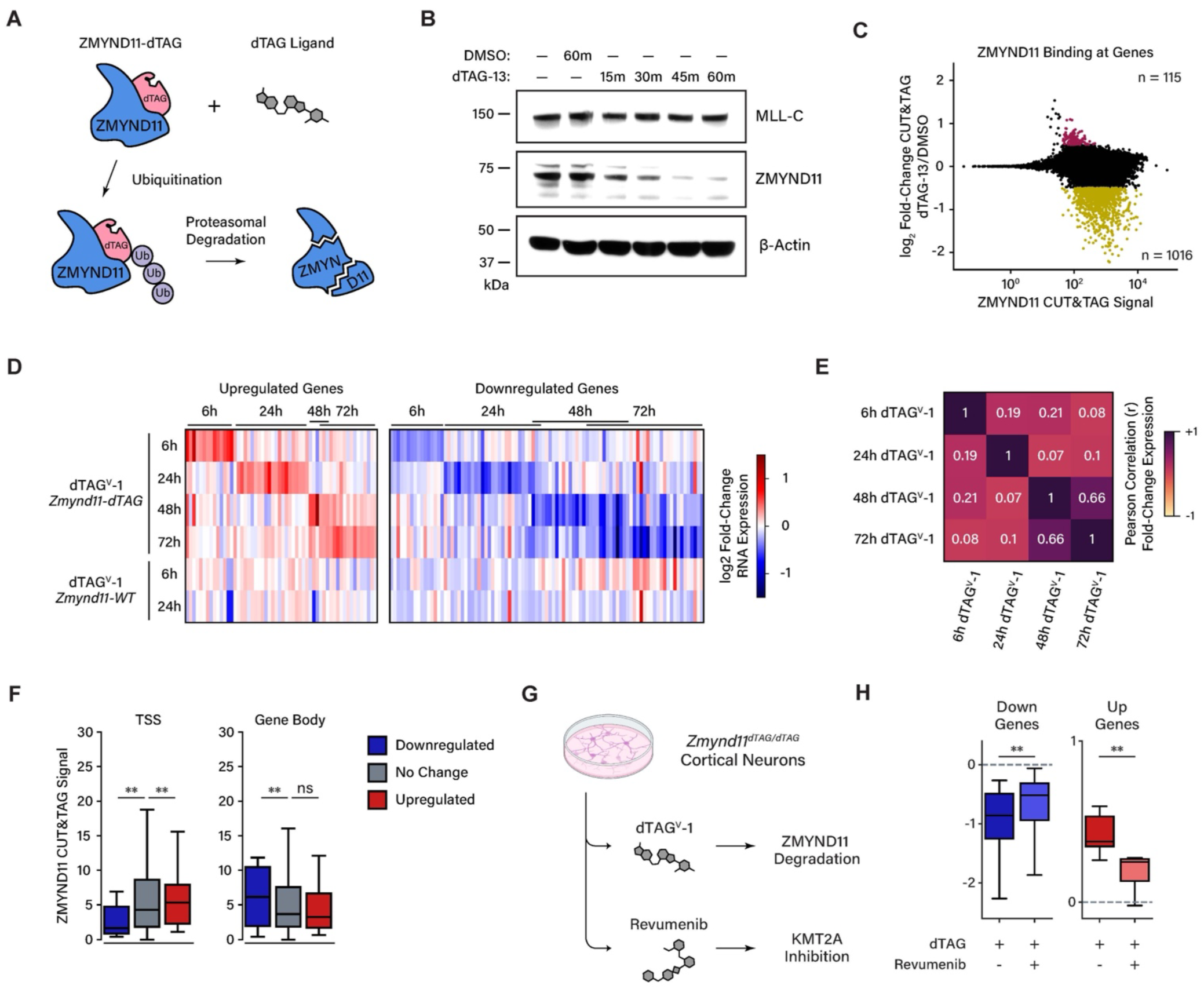
Rapid depletion of ZMYND11 and rescue by KMT2A inhibition. (a) Inducible degradation of FKBP12^F36V^-tagged ZMYND11 (*Zmynd11-dTAG*) protein using heterobifunctional dTAG small molecules. (b) Representative western blot showing ZMYND11, MLL-C, and β-Actin in *Zmynd11-dTAG* primary cortical neurons. Cells were treated with 500 nM dTAG-13 or DMSO vehicle. (c) Scatter plot showing ZMYND11 CUT&TAG signal in *Zmynd11-dTAG* cultured primary cortical neurons treated with 50 nM dTAG-13 for 3 hours, relative to DMSO vehicle control. Significant genes are highlighted (significance calculations by DESeq2, FDR-adjusted pval < 0.1, minimum log_2_ fold-change 0.5); n = 5 independent replicates. (d) Heatmap showing significant gene expression changes in *Zmynd11-dTAG* cultured primary cortical neurons treated with 500 nM dTAG^v^-1 for different durations from 6 to 72 hours. Plot shows DESeq2 log_2_ fold-change RNA expression, genes separated by significance in each time point of dTAG^v^-1 treatment tested. Significant genes filtered for FDR-adjusted pval < 0.1, minimum log_2_ fold-change 0.25. Total RNA-Seq, n = 4 independent replicates. (e) Heatmap showing Pearson correlation (r) of expression fold-change by gene in *Zmynd11-dTAG* primary cortical neurons treated with 500 nM dTAG^v^-1 for 6, 24, 48, or 72 hours; n = 4 independent RNA-seq replicates. (f) Enrichment of ZMYND11 CUT&TAG signal at the transcription start site (TSS) or gene body of genes grouped by significant upregulation, downregulation, or no change in expression in *Zmynd11-dTAG* primary cortical neurons upon treatment with 500 nM dTAG^v^-1 for 24 hours. ** = p < 0.01, p-values calculated using Wilcoxon rank-sum test; n = 6 independent CUT&TAG replicates. (g) Schematic showing experimental approach for treatment of *Zmynd11-dTAG* primary cortical neurons with dTAG^v^-1 and revumenib. (h) Change in expression of genes significantly misregulated in *Zmynd11-dTAG* primary cortical neurons with 500 nM dTAG^v^-1 administration for 24 hours relative to 500 nM dTAG^v^-1-NEG control, with and without co-administration of 2 µM revumenib. ** = p < 0.001, p-values calculated using Wilcoxon rank-sum test; n = 4 independent RNA-seq replicates.

Transcriptomic analysis of *Zmynd11-dTAG* neuronal cultures showed modest initial changes in gene expression upon dTAG treatment. By six hours, only a small number of genes significantly increased in expression, the majority of which were also up-regulated in dTAG-treated wild-type neurons, indicative of off-target effects of dTAG drugs **(Fig S10D)**. Beginning at 24 hours after drug treatment, a broader misregulated gene program began to emerge specific to *Zmynd11-dTAG* neurons, rather than wild-type neurons. The differentially expressed genes at 24 hours were transiently misregulated, giving way to a distinct later pattern of disrupted gene expression at 48 and 72 hours after drug treatment **(Fig 6D)**. The genes that were misregulated at 48 and 72 hours were highly correlated between those two time points and showed little correlation with genes misregulated at earlier time points **(Fig 6E)**. This suggests that direct targets of ZMYND11 may be the genes found to be misregulated by the 24 hour time point, and these genes may coordinate the regulation of a broader gene program whose expression is disrupted at later time points after ZMYND11 degradation. Notably, genes misregulated at six hours or 24 hours showed no enrichment for functions in neuronal development, whereas such functions emerged in gene ontology analysis in the 48 and 72 hour gene sets **(Fig S10E)**. Gene misregulation was stronger in neurons treated with dTAG^v^-1 than dTAG-13, suggesting that the former may be a more efficient degrader of ZMYND11, but differential gene expression compared to controls was correlated between both drug treatments **(Fig S10F)**.

It is noteworthy that while thousands of genes are significantly depleted of ZMYND11 binding upon ZMYND11 degradation, fewer than 200 genes were significantly misregulated across the 6-72 hour time points, further demonstrating that ZMYND11 binding alone is not sufficient to predict gene misregulation. We analyzed ZMYND11 CUT&TAG signal at the genes that are significantly misregulated upon ZMYND11 degradation and found that ZMYND11 binding is enriched at the start sites of up-regulated genes relative to genes which are down-regulated or unchanged. In agreement with our findings in *Zmynd11-cKO* brain tissue, this suggests that ZMYND11 binding at transcription start sites is predictive of gene repression by ZMYND11 **(Fig 6F)**.

If ZMYND11 inhibition of KMT2A is a primary mechanism by which ZMYND11 inhibits gene expression, we would expect that the effects of ZMYND11 degradation on gene expression would be reversed if KMT2A activity is inhibited. To test this hypothesis, we treated *Zmynd11-dTAG* neurons with dTAG^v^-1 for 24 hours in the presence or absence of revumenib, a KMT2A inhibitor approved for the treatment of *KMT2A*-rearranged leukemia^33^. Revumenib treatment significantly attenuated the transcriptional defects we observed upon ZMYND11 degradation, indicating that blocking KMT2A activity partially reverses the gene misregulation observed upon loss of ZMYND11 **(Fig 6G, S11A)**. While treatment with revumenib alone did not lead to statistically significant gene misregulation after 24 hours, the small changes in gene expression observed upon treatment with revumenib were anti-correlated with changes induced by ZMYND11 degradation, further supporting the interpretation that ZMYND11 and KMT2A regulate the expression of similar genes, but in opposite directions **(Fig S11B)**. Notably, only genes misregulated upon dTAG^v^-1 treatment that are specific to *Zmynd11-dTAG* neurons showed partial rescue by revumenib treatment, while genes misregulated by dTAG^v^-1 treatment in wild-type neurons (indicating off-target effects of the dTAG drug) were unaffected by KMT2A inhibition **(Fig S11C)**. These results suggest that the misregulation of gene expression observed upon by depletion of ZMYND11 is a consequence of increased KMT2A activity.

We next investigated the mechanism through which ZMYND11 is recruited to chromatin. While ZMYND11 is known to bind the histone mark H3K36me3, our earlier findings revealed that ZMYND11 binds to active transcription start sites and that this binding is particularly correlated with gene misregulation upon ZMYND11 loss, while H3K36me3 is restricted to active gene bodies rather than start sites. We performed CUT&TAG in *Zmynd11-dTAG* neurons treated with dTAG-13, and found that depletion of ZMYND11 does not alter KMT2A binding, indicating that KMT2A is not dependent on ZMYND11 for localization to chromatin **(Fig S11D)**. Conversely, treatment with revumenib strongly reduced KMT2A binding at transcription start sites, reflecting the key role of Menin in enabling chromatin binding of KMT2A. In revumenib-treated neurons, we also observed a concomitant decrease in CUT&TAG signal for ZMYND11 at start sites, whereas ZMYND11 binding at gene bodies was unaffected. These results suggest that KMT2A is required for binding of ZMYND11 to active start sites, while it is not required for ZMYND11 localization at gene bodies, reflecting the lack of KMT2A in such regions.

To test whether H3K36me3 is responsible for ZMYND11 binding at gene bodies, we treated neurons with EZM-0414, a selective inhibitor of SETD2 that prevents H3K36 trimethylation^49^. In these neurons we observed a strong depletion of H3K36me3 CUT&TAG signal across gene bodies, as well as a reduction in ZMYND11 signal, comparable in magnitude to the change observed upon degradation of ZMYND11 by dTAG-13. Unexpectedly, EZM-0414 also reduced ZMYND11 and KMT2A binding at transcription start sites, despite the lack of H3K36me3 at these sites. This raises the possibility of a mechanism through which SETD2 indirectly regulates KMT2A binding, thus causing a decrease in binding of ZMYND11 after SETD2 inhibition. Finally, degradation of ZMYND11 did not affect the deposition of H3K36me3, indicating that this mark is maintained independently of ZMYND11.

Taken together, these data support a model in which KMT2A recruits promoter-bound ZMYND11, whereas the H3K36me3 mark recruits ZMYND11 to gene bodies **(Fig 7A)**. By binding to KMT2A, ZMYND11 attenuates KMT2A-dependent gene activation. Critically, the ability of revumenib to rescue neuronal gene expression defects following acute ZMYND11 degradation suggests that inhibition of KMT2A is a core mechanism by which ZMYND11 maintains the balance of gene regulatory inputs in neurons, ultimately playing a key developmental role in the safeguarding of nervous system development.

**Figure 7.**
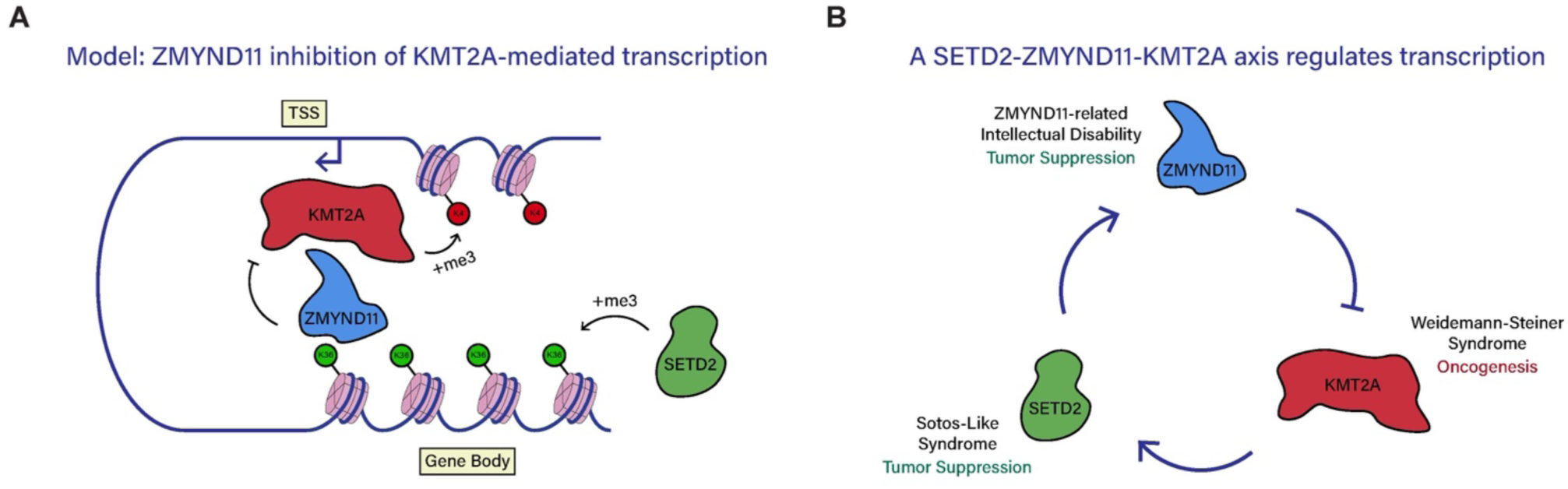
Model for ZMYND11 function. (a) Schematic illustrating proposed function of ZMYND11 as a reader of H3K36me3 across active gene bodies, and an inhibitor of KMT2A at active transcription start sites. ZMYND11 binds H3K36me3 across active gene bodies, the deposition of which is catalyzed by SETD2. ZMYND11 binds KMT2A at active transcription start sites, inhibiting its function and reducing gene activation. (b) Schematic showing a proposed transcriptional feedback loop consisting of ZMYND11, KMT2A, and SETD2, highlighting their complementary roles as regulators of neural development and cancer.

## Discussion

The number of genes linked to chromatinopathy disorders has increased with greater DNA sequencing in the clinical setting. Yet, despite these advances, the underlying mechanisms by which chromatin regulators contribute to brain development and dysfunction in many cases remain unclear. One explanation for the overlapping presentation of disorders linked to mutations of chromatin regulators is the convergence of such regulators upon common molecular pathways. Indeed, a significant number of chromatinopathy genes cluster into groups corresponding to complexes involved in mobilizing or modifying chromatin^4^. In this study, we uncover a link between ZMYND11 and the histone methyltransferase KMT2A, expanding the previous understanding of ZMYND11 as a reader of H3K36me3 and contextualizing its role in disease.

The molecular function of KMT2A in catalyzing H3K4me3 and promoting gene activation is shared across many tissues, and yet its disruption is felt most prominently in the brain, with loss-of-function mutations in *KMT2A* causing the chromatinopathy disorder Wiedemann-Steiner Syndrome (WSS)^29,30^. SETD2, the writer enzyme of H3K36me3, is also a chromatin regulator which has ubiquitous roles in gene regulation across cell types but plays a particularly key role in the brain, with haploinsufficiency of *SETD2* causing the neurodevelopmental syndrome Luscan-Lumish Syndrome (LLS)^50–52^. ZRSID, WSS, and LSS are thus all neurodevelopmental syndromes with significantly overlapping symptoms such as intellectual disability, delayed motor and language development, and seizures. Given the established importance of finely balancing levels of KMT2A activity for proper neuronal development, our findings offer a plausible explanation for the essential role of ZMYND11, as dysfunction of ZMYND11 leads to dysregulation of KMT2A.

We show the ability of the KMT2A inhibitor revumenib to partially rescue transcriptional dysfunction upon ZMYND11 degradation, supporting the conclusion that loss of ZMYND11 leads to increased activity of KMT2A. This finding also suggests that modulation of KMT2A activity should be investigated as a therapeutic approach for disorders caused by loss of ZMYND11 function, a question which can be explored in mouse models of ZMYND11 such as those described here. Revumenib inhibits the KMT2A cofactor Menin and has recently been approved as a first-in-class drug for the treatment of *KMT2A*-rearranged relapsed or refractory acute leukemia^34,53,54^. Revumenib and a second KMT2A-Menin inhibitor, ziftomenib, have also shown efficacy in the treatment of other cancers dependent on KMT2A function, such as *NPM1*-mutant leukemias^33,55,56^. Reflecting sustained interest in pharmacological approaches to target KMT2A, inhibitors have been developed to target different facets of KMT2A function, such as the cofactor WDR5, or catalytic activity of the KMT2A SET domain itself^57–60^. Given that we do not yet know the precise function of KMT2A inhibited by ZMYND11, inhibitors targeting different mechanisms of KMT2A function may provide further possibilities to compensate for ZMYND11 loss.

It is notable as well that KMT2A and ZMYND11 play opposite and complementary roles in cancer, with ZMYND11 functioning as a tumor suppressor while gain-of-function mutations in *KMT2A* drive oncogenesis. It should be considered that ZMYND11’s important roles in neuronal development and in cancer may both be mediated through inhibition of KMT2A. SETD2 is also a tumor suppressor, with *SETD2* mutations being found in a wide range of cancers, including in acute leukemia where *KMT2A*-rearranged leukemias more frequently harbor *SETD2* mutations than non-*KMT2A*-rearranged leukemias^61–63^. This raises the possibility of a complementary role for SETD2 and ZMYND11 in antagonizing cancer development, forming a coordinated axis in which SETD2-catalyzed H3K36me3 promotes ZMYND11 recruitment and inhibits KMT2A **(Fig 7B)**. In this model, ZMYND11 inhibition of KMT2A would reduce target gene transcription, thereby reducing H3K36me3 and ZMYND11 recruitment, constituting a transcriptional negative-feedback loop. Future work can examine untested aspects of this model, such as whether H3K36me3 regulates KMT2A binding at active start sites, and whether ZMYND11 interactions with KMT2A are facilitated by gene body H3K36me3.

Studies of gene regulation are often confounded by secondary effects that emerge rapidly upon disruption of gene regulatory networks. To identify direct target genes of ZMYND11, we employed a degron-tagged knock-in allele to enable rapid depletion of ZMYND11 protein in primary neurons. Results of these experiments revealed that the effects of ZMYND11 depletion on gene expression arise promptly but not immediately, with modest changes detectable by six hours, and greater changes at 24 hours. The small number of genes that are immediately upregulated include *Kdm5a*, a regulator of histone H3K4 demethylation, and *Epc1*, a component of the NuA4 histone acetyltransferase complex, suggesting how a small number of misregulated genes can potentially contribute to a broader cascade of misregulation. Notably, only beginning at 48 hours did we observe downregulation of genes linked to neuronal development such as *Astn2*, *Brinp1*, *Dcc*, and *Ostn*, which remained suppressed after 72 hours of dTAG^v^-1 treatment. These findings provide further evidence that ZMYND11 does not directly promote the expression of neuronal genes, but may enable their transcription by fine-tuning the levels of other developmental regulators to safeguard neuronal outcomes.

It has been broadly observed that mutation of genes linked to autism and chromatinopathy can cause either impaired neurogenesis or increased neurogenesis in stem cell models of neuronal differentiation, supporting a model in which proper brain development relies not just on the activation of particular developmental pathways, but on their precise balancing by a range of positive and negative inputs^64^. Consistent with our findings that ZMYND11 suppresses the activity of KMT2A, a recent study examining NDD-linked genes in a neuronal differentiation model reported opposite functions for ZMYND11 and KMT2A in neuronal differentiation. In these results, neural stem cells lacking ZMYND11 failed to differentiate into *TBR2^+^ FOXG1^+^* neurons, while cells lacking KMT2A showed increased efficiency of neuronal differentiation^24^. Along with these defects in differentiation, the authors observed up-regulation of genes linked to Wnt signaling in *ZMYND11* mutant neural stem cells. Similarly, we observed in *Zmynd11-cKO* brain tissue that genes linked to Wnt signaling were among the most highly upregulated. Wnt signaling has also been found to be upregulated upon loss of the chromatinopathy gene *Kdm5c*^65^, identifying the Wnt pathway as a possible locus of convergence in chromatinopathy. Interestingly, double-mutant mice bearing loss-of-function mutations in *Kdm5c* and *Kmt2a* rescue the neurodevelopmental phenotypes observed in mice bearing single mutations in either gene, suggesting that KDM5C and KMT2A play opposite roles in regulating brain development, possibly by exerting opposite effects on the activation of pathways regulating neural development such as Wnt signaling^66^. Taken together, these findings contribute to an emerging conceptual framework for the convergence of a wide range of chromatin mechanisms on signaling pathways regulating neuronal development.

Further aspects of ZMYND11 function, such as its interaction with the H3K36 dimethyltransferase and chromatinopathy-linked protein NSD1, remain promising new avenues for investigating the links between ZMYND11 and nervous system development, and ZMYND11’s role as a potential mediator of crosstalk between diverse chromatin marks. The resources we have generated in this study will enable further investigation into the function of ZMYND11, both at the level of its molecular function and its role in disease. The similarity of behavioral disturbances in *Zmynd11-cKO* mice and those in ZRSID suggest that these mice may serve as a useful model for studying underlying mechanisms of the human disorder and possible therapeutic interventions. Notably, our observation that the KMT2A inhibitor revumenib partially reverses gene misregulation caused by ZMYND11 depletion suggests that revumenib and other KMT2A inhibitors should be assessed in mouse models to determine whether they hold potential for ameliorating developmental or behavioral consequences of ZMYND11 disruption.

## Supporting information

supp_figures_with_legends

## Author Contributions

A.W.G. and M.E.G. conceived the study, designed experiments, and wrote the manuscript. A.W.G. performed the majority of experiments in the study. X.S.W. performed *in vitro* immunoprecipitation experiments and immunoblotting, and contributed to the writing of the manuscript. E.C.G. contributed to the writing of the manuscript. J.E.R. and M.D. assisted with histology and biochemical assays. I.R.V., L.T., S.K., and K.A.F. performed and analyzed behavioral experiments.

## Acknowledgements

The authors would like to acknowledge members of the Greenberg lab members for valuable discussion and input. We thank Pingping Zhang for invaluable assistance with mouse colonies. Protein mass spectrometry was performed by the Taplin Mass Spectrometry Facility and the Thermo Fisher Scientific Center for Multiplexed Proteomics at Harvard Medical School. Additional behavioral testing was performed by the IDDRC Animal Behavior and Physiology Core at Boston Children’s Hospital, funded by NIH/NICHD P50 HD105351. The Greenberg Laboratory is supported by the Allen Discovery Center Program, a Paul G. Allen Frontiers Group advised program of the Paul G. Allen Family Foundation, and the Tang-Yang Autism Center at Harvard Medical School. A.W.G. was supported by an NSERC Postgraduate Doctoral Scholarship and a Stuart H.Q. & Victoria Quan Fellowship. X.S.W was supported by the Hock E. Tan and K. Lisa Yang Center for Autism Research. M.E.G. was supported by funding from NINDS R01 NS115965. The funders had no role in study design, data collection and analysis, or preparation and publication of the manuscript.

## Supplementary Figures

Supplementary Figure 1: Germline disruption of ZMYND11 impacts mouse development in homozygotes but not heterozygotes. Related to Figure 1.

Supplementary Figure 2: Behavioral testing of *Zmynd11-cKO* mice. Related to Figure 1.

Supplementary Figure 3: Analysis of gene expression in *Zmynd11-cKO*. Related to Figure 2.

Supplementary Figure 4: Analysis of alternative splicing in *Zmynd11-cKO*. Related to Figure 2.

Supplementary Figure 5: Analysis of ZMYND11 genomic binding patterns. Related to Figure 3.

Supplementary Figure 6: Generation and validation of *Zmynd11-FH* mouse line. Related to Figure 4.

Supplementary Figure 7: Association of ZMYND11 with KMT2A. Related to Figure 4.

Supplementary Figure 8: Screening interactions of ZMYND11 variants associated with neurodevelopmental disorders. Related to Figure 5.

Supplementary Figure 9: Repression of reporter gene expression by ZMYND11. Related to Figure 5.

Supplementary Figure 10: Generation and characterization of *Zmynd11-dTAG* mouse line. Related to Figure 6.

Supplementary Figure 11: Interrogation of ZMYND11 function and binding using inhibitors of KMT2A and SETD2. Related to Figure 6.

## Experimental Models

### Mouse Models

Animals were housed and handled in accordance with protocols established by the Harvard Medical School Institutional Animal Care and Use Committee and in accordance with federal guidelines. The following mouse lines were used: wild-type C57/BL6 (Jackson Labs Stock 000664), *EIIa-Cre* (Jackson Labs Stock 003724), *Baf53b-Cre* (Jackson Labs Stock 027826), *Zmynd11-fl* (this study), *Zmynd11-KO* (this study), *Zmynd11-FH* (this study), *Zmynd11-dTAG* (this study). Male and female mice were used in equal proportions. Control mice were age- and sex-matched, and were littermates whenever possible.

### Generation of *Zmynd11* Mouse Lines

Zygote injections were performed by the Harvard Genome Modification Facility under the supervision of Dr. Lin Wu. Guide RNA sequences were designed to target regions flanking Exon 7 (*Zmynd11-fl*) or the beginning of the *Zmynd11* coding sequence (*Zmynd11-FH*, *Zmynd11-dTAG*) and chemically synthesized by Synthego as a chimeric single-guide RNA (sgRNA). Repair templates were designed with homology arms of 300 bp on either side of the sequence to be edited and synthesized as a single-stranded DNA fragment by Integrated DNA Technologies (IDT). Guide RNA, ssDNA repair template, and Cas9 protein were injected in C57/BL6 zygotes by the Harvard Genome Modification Facility. Offspring were screened by PCR and Sanger sequencing to identify F0 founders. F1 offspring were further screened by PCR and Sanger sequencing to validate genome editing.

### Mouse Neuron Cultures

Embryonic mouse cortex tissue was dissected from embryonic day 16.5 (E16.5) mouse litters from 6-8 embryos of mixed sex and pooled for each replicate culture. Cells were dissociated using a DNAse/Papain mixture for 10 minutes at 37°C with gentle agitation every 3 minutes. Trypsin was inhibited with ovomucoid inhibitor. Digested tissue was gently washed with dissection media and triturated by gentle pipetting with a P1000 tip. Dissociated cells were passed through a 40 μm filter and counted. Neurons were plated on cell culture dishes coated overnight with a solution of poly-D-lysine (20 μg/mL) and laminin (4 μg/mL). Neurons with grown in Neurobasal media (Gibco) supplemented with B-27 Supplement (Thermo Fisher), GlutaMAX (Thermo Fisher), 50 U/mL penicillin, and 50 U/mL streptomycin. Neurons were cultured in an incubator at 37°C with a CO2 concentration of 5% for 7 days before use in experiments.

## Experimental Procedures

### Golgi Staining

Samples were prepared for Golgi staining using the FD Rapid GolgiStain Kit (FD NeuroTechnologies) in accordance with manufacturer’s protocols. Brains were dissected from 6-week-old mice, washed in Milli-Q water, and impregnated for two weeks in a mixture of Golgi Solutions A + B. Brains were then transferred into Golgi Solution C for 48 hours, with the solution changed after the first day. Brains were frozen by immersion in a cryogenic bath of 1-butanol and dry ice and sectioned on a cryostat. Sections (100 μm) were mounted on gelatin-coated glass slides and dried at room temperature. Sections were washed in Milli-Q water and stained in a mixture of Golgi Solutions D+E, then washed again in water and dehydrated in subsequent baths of 50%, 75%, 95%, and 100% ethanol. Sections were then cleared in xylenes, and a coverslip was mounted with Eukitt mounting medium (Sigma Aldrich).

### Imaging of Golgi Stained Sections

Golgi-stained sections were imaged on a Zeiss Axioskop microscope with brightfield illumination, using Neurolucida software (MBF Bioscience) for live tracing of the dendritic arbor of individual Layer 2/3 pyramidal neurons within primary somatosensory cortex. Imaging was performed blinded to the genotype of each sample. Tracing was carried out at 63X magnification, and summary images were acquired at 20X magnification at 1600x1200 pixel resolution.

### Housing for Behavioral Testing

Behavioral testing was carried out with 2-3-month-old mice. For contextual fear conditioning, object recognition, and social recognition tasks, mice were kept in type III closed-lid standard cages (Tecniplast green line) in temperature-controlled rooms on a constant 12-hour light-dark cycle, and all experiments were conducted at approximately the same time of the light cycle (9am-1pm). No handling, except for routine cage changes once per week were performed on animals prior to the behavioral experiments. Before behavioral tests, animals were acclimatized to the behavior room for a minimum of 5 days. All experimental arenas/contexts were placed within a soundproof box equipped with camera and illumination systems to ensure consistent conditions throughout all experiments without unforeseen disturbance or variations from the surroundings. For behavioral experiments, animals were carefully taken out of their home cage and carefully transported to the testing arena with minimal stress to the animals. After the behavioral procedure, animals were allowed to voluntarily climb on the lid of their home cage and transferred back to their home cage.

For marble burying, rotarod, and elevated plus maze tasks, mice were kept in Optimice cages in temperature-controlled rooms on a constant 12-hour light-dark cycle. Before behavioral tests, animals were acclimatized to the behavior room on the day of testing. All experimental arenas/contexts were kept within a quiet environment with limited disturbances.

### Contextual Fear Conditioning

Five days prior to testing, mice were transferred to a dedicated behavior assay room and single housed with a 12 hour light/dark cycle. Contextual fear conditioning training was performed within a Coulbourn Instruments Habitest Modular Fear Conditioning Chamber (19cm x 18cm x 33cm) housed inside of a sound isolation cubicle. Context consisted of a plexiglass arena with a floor composed of metal rods and a 2% acetic acid odorant supplied via a petri dish within the waste collection area under the floor. Chambers were cleaned with 70% ethanol before each animal was tested. In contextual fear conditioning acquisition sessions, mice were introduced to the arena and recorded for a total of 360 seconds. Mice received five 1-second, 0.8 milliamp shocks at 30 second intervals beginning at 180 seconds after introduction to the chamber and concluding at 300 seconds. Mice were returned to their home cage 60 seconds after the final shock was administered and undisturbed for 24 hours. The following day, recall and extinction testing was performed by reintroducing mice to the context of the chamber for a total of 30 minutes without any further shock administration. Freezing behavior was identified within videos using FreezeFrame software with a freezing detection threshold of 1.0 and a minimum bout duration of .25 seconds. Freezing rates were calculated as percentage of time spent freezing within time bins representing acquisition (final 60 seconds post final shock of acquisition session), recall (first 5 minutes of recall session), and extinction (final 5 minutes of recall session).

### Object and Social Recognition Tasks

Object and social recognition tasks were performed as previously described^67^. On Day 1 of testing, mice explored two identical objects (50 mL Falcon tubes) placed in a 30x50 cm rectangular arena for 10 minutes. Mice were returned to their home cage immediately after training, and were tested for familiar object recognition after 1 hour (short-term memory, STM) and after 24 hours (long-term memory, LTM) for a period of 5 minutes. For the object recognition test, one of the two objects was replaced with a novel object (a tower built from Lego bricks). Following the long-term object recognition test on Day 2, mice were allowed to recover in their home cage for 1 hour before being returned to the arena and presented with an intruder animal (5 weeks old; not previous littermate) shielded inside a cage for 10 minutes.

To avoid discrimination of the objects based on odor, both the arena and the objects were thoroughly wiped with 70% ethanol before and after each trial. Only direct sniffing contact towards the objects was counted as interaction. To automatically quantify the behavioral sniffing responses towards objects and social intruders, we made use of the previously developed markerless pose estimation tool DeepLabCut^68^ (DLC). Briefly, DLC was used to train a single animal model of 8 keypoints (body parts: ear_left, ear_right, nose, center, lateral_right, lateral_left, tail_base, tail_tip) within our behavioral arena from 15 videos. Keypoints were inspected, and additional frames were manually annotated and corrected. The model was further evaluated with example videos left out from the initial model training. Training, evaluation and analysis configuration: TrainingFraction: - 0.95; iteration: 5; default_net_type: resnet_50; default_augmenter: default; snapshotindex: -1; batch_size: 8. Filtered pose estimates generated in DLC were then used to build a classifier for investigation (independent of object location) in SimBA^69^. Frames (92,902 total; 9832 frames positive for object or social exploration) from 17 videos were manually annotated to generate training data for the investigation classifier (independent of specific objects). We operationally defined investigation as active sniffing, interaction, or rearing on objects or social intruder cup. The SimBA GUI was used to create a random forest classifier with the following hyperparameters: criterion=gini, minimum sample leaf=1, number of estimators=2000, train/test size = 0.2, and maximum features=sqrt. The model performance was evaluated with a precision-recall curve and validated with an example video excluded from the initial model training. The probability threshold was set at 0.08, which minimized the number of false negative frames and false positive frames in two validation videos. In order to incorporate the object location into the investigation classifier, rectangular regions of interest (ROIs) around each object or social intruder cup were manually drawn, from which the hypotenuse was calculated. The distance of the nose to each ROI center was then calculated over time. The nose distance was used to define a spatial filter for the classifier. Specifically, the spatial filter was used to identify frames where the mouse was close to specific objects and was defined when the nose distance to ROI center was less than the hypotenuse of each object plus a 22 pixel (4 cm) approach border. The behavioral classifier in addition to the spatial filter identified frames where per-object investigation occurred, and the accuracy of this final model (behavioral classifier model plus spatial filter) was evaluated for a single video and represented in a confusion matrix. This final model was used to calculate time spent investigating the novel (or social intruder) or conditioned object. The preference index was calculated as (investigation time of novel object – investigation time of conditioned object)/(total object investigation time), where positive values indicate more time investigating the novel object. Investigation bouts were defined as per object interaction time separated by more than 2 frames.

### Marble Burying Test

Mice were introduced into a clean cage containing 3-4 cm of fresh, Biofresh Performance Bedding. Following a 10-minute habituation period, the mice were temporarily transferred to a secondary cage while the bedding was leveled, and 20 glass marbles (arranged in a 4 x 5 grid) were placed on top of the bedding. Subsequently, each mouse was returned to the cage for 10 minutes, and the number of marbles buried to a depth covering 50% of their surface area was recorded. To eliminate any residual scent from previous mice, the bedding was replaced for each subsequent mouse, and the marbles were thoroughly cleaned by soaking them in Peroxigard and subsequently dried within the bedding material between successive tests.

### Rotarod Test

To acclimate to the movement of the rotarod, mice were placed on the rotarod (IITC Life Science Inc., USA) set to revolve at 5 rpm for 5 continuous minutes. Mice were placed back on the rod immediately after falling. On the second day, mice were placed on the rotarod, and after a 10 second acclimation period with the rod revolving at 4 rpm, it then began to accelerate at 0.1 rpm/sec. When the mouse fell from the rod, the latency to fall was recorded. This test trial was repeated three times with a break of 5 minutes between trials. The average time across trials was calculated as the final result for the mice. Testing was repeated for three consecutive days.

### Elevated Plus Maze

Mice were habituated in the testing room for least 30 min prior to start of testing. Mice were placed in an elevated plus maze (EPM), consisting of two open arms and two closed arms. All four arms were elevated 1 meter from the floor, with the drop-off detectable only in the open arms. Testing was performed according to previously described procedures using a mouse EPM (model ENV-560A, Med Associates, St. Albans, VT)^70^. The EPM contained two open arms (35.5 cm x 6 cm) and two closed arms (35.5 cm x 6 cm) radiating from a central area (6 cm x 6 cm). A 0.5 cm high lip surrounded the edges of the open arms, whereas the closed arms were surrounded by 20 cm high walls. The EPM was cleaned with Peroxigard before the beginning of the first test session and after each subject mouse was tested, with sufficient time for the ethanol odor to dissipate before the start of the next test session. Room illumination was ∼30 lux. To begin the test, the mouse was placed in the central area facing the open arm. The mouse was allowed to freely explore for 5 minutes. Trials were recorded on video and scored with EthoVision XT video tracking software (version 15.0, Noldus Information Technologies, Leesburg, VA).

### Nuclear Isolation from Brain Tissue

Brain tissue (whole cortex for CUT&RUN and RNA, whole brain including cerebellum for immunoprecipitation) was dissected from 6-week-old mice. Tissue was placed in homogenization buffer (250 mM sucrose, 25 mM KCl, 5 mM MgCl2, 20 mM Tricine-KOH pH 7.8) and dissociated with a dounce homogenizer. After dissociation with the tight pestle, IGEPAL CA-630 was added to a final concentration of 0.32% and five more strokes were applied. Lysates were combined with an equal volume of OptiPrep (Sigma-Aldrich) mixed with diluent buffer (150 mM KCl, 30 mM MgCl2, 120 mM Tricine-KOH pH 7.8) for a final concentration of 25% iodixanol. In a clear ultracentrifuge tube, a gradient of 30% iodixanol was layered on top of 40% iodixanol. The 25% iodixanol lysate mixture was carefully pipetted on top of the 30/40% iodixanol gradient and centrifuged at 7600 RPM for 18 minutes in an SW-41 rotor. Nuclei were isolated by pipetting at the gradient interface and diluted with one part buffer NE1 (10 mM KCl, 3 mM MgCl2, 0.1% Triton X-100, 20 mM HEPES pH 7.9).

### Stereotactically Guided Surgery

All surgeries were carried out in accordance with protocols approved by the Harvard Medical School Institutional Animal Care and Use Committee and in accordance with federal guidelines. Mice were anesthetized by isoflurane inhalation (3-4% induction, 1-2% maintenance) within a stereotaxic frame (Kopf) and maintained at 37°C on a heating pad. Fur over the scalp was removed using a depilatory containing potassium thioglycolate and calcium hydroxide and sterilized with three alternating washes of betadine and 70% ethanol. A burr hole was drilled through the skull above the primary visual cortex (V1), and a glass pipette filled with adeno-associated virus (AAV) was lowered into V1 (coordinates: AP -3.5, ML +/- 2.5, DV -0.60). AAV (1 μL, diluted to 1.0 × 10^12^ genome copies per mL) was injected at a rate of 150 nL/min. The pipette left in place for 1 minute to enable diffusion of virus, withdrawn to a depth of -0.40, and then left for 1 minute longer before fully withdrawing. All animals were given postoperative analgesic (buprenophine slow-release formulation, 1 mg/kg) and monitored post-operatively. Mice were injected at an age of four weeks.

### Lentiviral Infection

Lentiviral plasmids were delivered into HEK293T cells by BBS/CaCl2 transfection. 16-20 hours after transfection, cells were assessed for fluorescence and changed into fresh culture media. After 24 hours, viral media was collected and filtered using a 0.45 μM filter. Virus was pelleted by ultracentrifugation at 25,000 rpm for 90 minutes at 4°C, and then resuspended by incubation in PBS overnight at 4°C. Concentrated virus was then stored at -80°C. Cultured mouse neurons were infected by adding concentrated lentivirus to culture media at day 2 *in vitro* (DIV2).

### RNA Isolation and RNA-Seq Library Preparation

RNA was isolated from tissue or nuclei homogenized and stored in Trizol (Invitrogen) using chloroform extraction and purified using a RNeasy Micro Kit (QIAGEN) with on-column DNase digestion. RNA libraries were generated with the NEBNext Ultra Directional II Library Prep Kit with rRNA Depletion (NEB). Other RNA libraries were prepared using the NEBNext UltraExpress Library Prep Kit with rRNA Depletion Kit (NEB). Libraries were sequenced on an Illumina NextSeq 500 with 80x80 bp paired-end reads, or on an Illumina Novaseq X with 150x150 bp paired-end reads. All RNA libraries for a given experimental sample were prepared simultaneously with paired controls.

### Precision Run-On Sequencing (PRO-Seq)

Cultured neurons were washed three times in cold PBS and then scraped in 1 mL Permeabilization Buffer (10 mM Tris-HCl pH 8.0, 10 mM KCl, 250 mM sucrose, 5 mM MgCl2, 0.1% IGEPAL CA-630, 0.5 mM DTT, 10% glycerol). Cells were centrifuged at 700*g* at 4°C for 5 minutes, then resuspended in 100 μL Wash Buffer (10 mM Tris-HCl pH 8.0, 10 mM KCl, 250 mM sucrose, 5 mM MgCl2, 0.5 mM DTT, 10% glycerol). Permeabilization Buffer (900 μL) was added and cells mixed gently by pipetting. Cells were centrifuged at 700*g* at 4°C for 5 minutes, then resuspended in 200 μL Freezing Buffer (50 mM Tris-HCl pH 8.0, 5 mM MgCl2, 0.5 mM DTT, 40% glycerol). Cells were centrifuged at 700*g* at 4°C for 5 minutes, then resuspended in 50 μL Freezing Buffer. Cells were snap-frozen in liquid N2 and stored at -80°C.

Frozen nuclei were thawed on ice and combined with 50 μL 2X Run-on Buffer (10 mM Tris-HCl pH 8.0, 5 mM MgCl2, 300 mM KCl, 40 μM Biotin-11-ATP, 40 μM Biotin-11-GTP, 40 μM Biotin-11-CTP, 40 μM Biotin-11-UTP, 1% Sarksosyl, 1 mM DTT, 0.8 units/μL SUPERase In RNase Inhibitor) and incubated at 37°C for 5 minutes. The reaction was quenched with 500 μL Trizol LS, and RNA was extracted with 130 μL chloroform and centrifuged at 17,000*g* for 5 minutes at 4°C. The aqueous layer containing RNA was combined with 1 mL ice-cold ethanol and 2.5 μL GlycoBlue (Invitrogen) and centrifuged at 17,000*g* for 15 minutes at 4°C. The RNA pellet was washed in 75% ethanol, air-dried, and resuspended in 20 μL nuclease-free water.

RNA was hydrolyzed by adding 5 μL 1 N NaOH and incubating for 4 minutes at 4°C, then neutralized with 30 μL 1 M Tris-HCl pH 7.0. Hydrolyzed RNA was run on a Micro Bio-Spin P-30 Column (Bio-Rad) to remove excess salts. Biotinylated RNA was purified with magnetic streptavidin beads (NEB) at room temperature for 20 minutes and washed 2 times each with 500 μL High-Salt (2 M NaCl, 50 mM Tris-HCl pH 7.4, 0.5% Triton X-100, 4 units/mL SUPERase In RNase Inhibitor), Binding (300 mM NaCl, 10 mM Tris-HCl pH 7.4, 0.1% Triton X-100, 4 units/mL SUPERase In RNase Inhibitor), and Low-Salt (5 mM Tris-HCl pH 7.4, 0.1% Triton X- 100, 4 units/mL SUPERase In RNase Inhibitor) wash buffers. Beads were resuspended in 300 μL Trizol and combined with 60 μL chloroform, and RNA was purified as above.

The dried RNA pellet was resuspended in 4 μL of 12.5 μM Reverse 3’ RNA adaptor and used to make to a T4 RNA Ligase I reaction mix (5 μM Reverse 3’ RNA Adaptor, 1X T4 RNA Ligase I Buffer, 1 mM ATP, 10 units T4 RNA Ligase I, 10% PEG 8000, 2 units/μL SUPERase In RNase Inhibitor). The reaction was incubated overnight at 20°C. The ligated RNA was purified with magnetic streptavidin beads, washed, and extracted with Trizol as above. The RNA was treated with Rpph to remove the 5’ cap (1X NEBuffer 2, 10 units RppH, 0.2 units/μL SUPERase In RNase Inhibitor). The reaction was incubated for 1 hour at 37°C. A hydroxyl repair reaction mix (1X PNK Buffer, 1 mM ATP, 25 units PNK, 0.2 units/μL SUPERase In RNase Inhibitor) was added directly to the Rpph reaction and incubated for an additional hour at 37°C. The RNA was extracted with Trizol as above. The dried RNA pellet was resuspended in 4 μL of 12.5 μM Reverse 5’ RNA adaptor and used to make to a T4 RNA Ligase I reaction mix (5 μM Reverse 5’ RNA Adaptor, 1X T4 RNA Ligase I Buffer, 1 mM ATP, 10 units T4 RNA Ligase I, 10% PEG 8000, 2 units/μL SUPERase In RNase Inhibitor). The reaction was incubated overnight at 20°C. The ligated RNA was purified with magnetic streptavidin beads, washed, and extracted with Trizol as above. The resulting RNA library was reverse transcribed using SuperScript III Reverse Transcriptase (Invitrogen). The cDNA libraries were amplified using Q5 DNA Polymerase (NEB) for 14 cycles or more as determined by a test amplification with serially diluted cDNA. Amplified libraries were purified using Agencourt RNAClean XP beads. Libraries were sequenced on an Illumina Nextseq 500 with 75 bp single-end reads. All PRO-Seq libraries for a given experimental sample were prepared simultaneously with paired controls.

#### CUT&RUN

CUT&RUN experiments were performed on nuclei isolated from brain tissue as described above. 250,000 nuclei per condition were resuspended in a final volume of 1 mL of CUT&RUN wash buffer (150 mM NaCl, 0.2% Tween-20, 20 mM HEPES pH 7.5, 1 mg/mL BSA, 10 mM sodium butyrate, 0.5 mM spermidine, 1X cOmplete EDTA-free protease inhibitor). Magnetic concanavalin-A (ConA) beads (Bangs Laboratories) were washed with CUT&RUN binding buffer (10 mM KCl, 1 mM CaCl2, 1 mM MnCl2, 20 mM HEPES-KOH pH 7.9) and added to each sample to bind nuclei. Nuclei bound to beads were resuspended in CUT&RUN antibody buffer (wash buffer + 0.1% Triton X-100 + 2 mM EDTA) and incubated overnight with 1 μL of antibody at 4°C. After overnight incubation, nuclei on beads were washed with antibody buffer and resuspended in CUT&RUN triton wash buffer (wash buffer + 0.1% Triton X-100). Protein-A-MNase (pA-MNase) was added to a final concentration of 700 ng/mL and samples incubated for 1 hour at 4°C. Samples were washed three times with triton wash buffer, then resuspended in 100 μL triton wash buffer. CaCl2 was added at a final concentration of 3 mM to activate MNase enzyme activity and incubated for 1 hour at 4°C. To stop the reaction, 100 μL of 2X STOP buffer (340 mM NaCl, 20 mM EDTA, 4 mM EGTA, 0.04% Triton X-100, 0.1 μg/mL RNase A) was added to each reaction and incubated for 20 min at 37°C. Beads were captured on a magnet, and supernatant containing CUT&RUN DNA was transferred to another tube. Two microliters of 10% SDS and 2 μL of 20 mg/mL Proteinase K were added to each sample and incubated for 1 hour at 65°C, shaking at 600 RPM. DNA was extracted using Phenol-Chloroform Isoamyl Alcohol, with ethanol precipitation of the aqueous layer. CUT&RUN libraries were prepared using the ThruPLEX DNA-Seq Kit (Takara). Libraries were sequenced on an Illumina Nextseq 500 with 40x40 bp paired-end reads. All CUT&RUN libraries for a given experimental sample were prepared simultaneously with paired controls.

#### CUT&TAG

Cultured neurons were washed once in cold PBS and then scraped in 1 mL NE1 buffer (10 mM KCl, 3 mM MgCl2, 0.1% Triton X-100, 20 mM HEPES pH 7.9). Nuclei were centrifuged at 700*g* at 4°C for 5 minutes, then resuspended in μL NE1. 250,000 nuclei per condition were aliquoted as for CUT&RUN above, bound to magnetic ConA beads, and incubated with primary antibody overnight at 4°C. After overnight incubation, nuclei on beads were resuspended in antibody buffer with 1 μL guinea pig anti-rabbit secondary antibody (Fisher) and incubated at RT for 30 minutes. Nuclei were then washed twice with 500 μL wash buffer. pA-Tn5 adaptor complex was diluted in WB300 (wash buffer + NaCl to 300 mM) with a ratio of 2.5 μL pA-Tn5 complex (0.5 mg/mL) in 100 μL WB300. Nuclei were then resuspended in WB300+pA-Tn5 complex and incubated for 1 hour at RT. Nuclei were washed twice with 500 μL WB300, and then resuspended in 300 μL tagmentation buffer (WB300 + 10 mM MgCl2) and incubated with agitation for 1 hour at 37°C. Tagmentation was halted by adding 10 μL 0.5M EDTA, 3 μL 10% SDS, and 2 μL 20 mg/mL Proteinase K (Thermo Fisher) and incubating nuclei for 1 hour at 50°C. DNA was extracted by adding 300 μL phenol-chloroform-isoamyl alcohol (Invitrogen) and taking the aqueous phase. The aqueous phase was extracted with 300 μL chloroform and added to a fresh tube, then precipitated by adding 750 μL 100% ethanol and 2 μL GlycoBlue (Invitrogen). This mixture was incubated for 1 hour at -20°C, then centrifuged at 17,000*g* for 10 minutes at 4°C. The DNA pellet was washed with 75% ethanol, then dried for 5 minutes at RT. The DNA pellet was resuspended in 20 μL ultrapure water and stored at -20°C. Libraries were amplified using KAPA HiFi non-hot-start DNA polymerase (Roche). Libraries were sequenced on an Illumina Nextseq 500 with 40x40 bp paired-end reads, or on an Illumina Novaseq X with 150x150 bp paired-end reads. All CUT&TAG libraries for a given experimental sample were prepared simultaneously with paired controls.

### Immunoprecipitation of ZMYND11 from Brain

Immunoprecipitation was performed using nuclei isolated from whole brain as described above. Nuclei diluted in buffer NE1 were centrifuged at 700*g* at 4°C for 2 minutes and resuspended in 400 μL buffer NE1. NaCl was added to a final concentration of 420 mM, and nuclei were rotated at 4°C for 20 minutes to dissociate chromatin. Nuclei were then centrifuged at 12,000*g* at 4°C for 10 minutes, causing insoluble chromatin to form a gelatinous pellet at the bottom of the tube. The soluble fraction was set aside, and chromatin was digested using micrococcal nuclease (200 μL MNase buffer (NEB), 5 μL MNase enzyme (NEB), 1% cOmplete EDTA-free protease inhibitor tablet (Sigma-Aldrich), 1 mM DTT). For MNase digestion, chromatin mixture was incubated at 37°C while shaking at 1000 RPM for 5 minutes. The solubilized chromatin fraction was combined with the soluble nucleoplasm fraction and diluted with NE1 to a final concentration of 150 mM NaCl. Inputs were pre-cleared by incubation with 50 μL of Protein A/G Magnetic Agarose (Thermo Fisher) for 1 hour at 4°C. Pre-cleared inputs were incubated with 30 μL anti-DYKDDDK Magnetic Agarose (Pierce) overnight at 4°C. Beads were washed three times with buffer NE1+250 mM NaCl. Samples were eluted from beads using 3XFLAG peptide at 500 μg/mL in buffer NE1+250 mM NaCl, shaking at 1000 RPM for 10 minutes at room temperature. Samples were precipitated with TCA and analyzed by the Taplin Mass Spectrometry Facility (Harvard Medical School) using protein LC/MS-MS. All brain immunoprecipitation experiments were performed using two *Zmynd11^FH/FH^* and two *Zmynd11^WT/WT^* littermate controls per experiment.

### Immunoprecipitation of ZMYND11 from HEK293T Cells

Plasmids expressing FLAG-HA-tagged ZMYND11 constructs or GFP (pLJM1 with UbC promoter) were expressed in HEK293T cells by CaCl2-BBS transfection (10 μg DNA, 10 million cells). Tagged ZMYND11 constructs were expressed as a bicistronic expression construct with eGFP-P2A to enable visualization of transfection efficiency. Cells were washed once with cold PBS and scraped into 2 mL cold PBS. Cells were centrifuged at 700*g* for 5 min at 4°C and resuspended in 5 mL buffer NE1. Cells were rotated for 10 min at 4 °C to ensure lysis, then centrifuged at 2000*g* for 5 min at 4°C. The cell pellet was resuspended in 200 μL (approximately one pellet volume) of buffer NE1 + 2 μL Benzonase nuclease (Sigma Aldrich). To digest DNA, cells were rotated at for 5 min at room temperature and then for 1 hour at 4°C. NaCl was added to a concentration of 420 mM to inactivate benzonase, and lysates rotated for 20 min at 4°C. NE1 was added to dilute the final concentration of NaCl to 250 mM and lysates spun at 12,000*g* for 20 min at 4°C to pellet insoluble material. The remaining soluble lysate was used as input for immunoprecipitation. Samples were pre-cleared, immunoprecipitated, washed, and eluted as for brain anti-FLAG IP, above. Samples were precipitated with TCA and analyzed by the Thermo Fisher Center for Multiplex Proteomics (Harvard Medical School) using tandem mass tag multiplex protein mass spectrometry (TMT-MS). Two independent immunoprecipitation experiments per condition were labeled and pooled for analysis on a 10-plex TMT-MS run.

### Immunoblotting

Extracts from brain tissue were resolved on 4-12% Bis-Tris gels (Thermo Fisher) and transferred to nitrocellulose membranes. Membranes were blocked for 1 hour at room temperature with 4% BSA in TBS-T, incubated with primary antibodies overnight at 4°C, washed 3 times with TBS-T, incubated with secondary antibody for 45 minutes at room temperature, then washed 3 times with TBS-T. Western blots incubated with fluorescent secondary antibodies were imaged on a Li-Cor Odyssey. Western blots incubated with HRP-coupled secondary antibodies were imaged by chemiluminescence and film exposure.

### Generation and Selection of Reporter K562 Cell Lines

HEK293T cells were transfected with lentiviral plasmids by CaCl2-BBS transfection to produce virus expressing rTetR-effector fusion constructs. Culture medium was changed 24 hours post-transfection, and viral media collected 48- and 72-hours post-transfection and sterile filtered through a 0.45 μm low protein-binding filter. K562 cells previously engineered to harbor a Citrine fluorescent reporter locus regulated by Tet operator sites (TetO) were transduced by resuspending cells in lentiviral media and centrifuging at 1000*g* for 1 hour at 32°C ^46^. Cells were resuspended in RPMI growth medium and incubated for three days before selection with 10 μg/mL Blasticidin S (Gibco), passaging every 2-3 days until only successfully transduced RFP-expressing cells remained. K562 reporter cells were treated with 1 μg/mL Doxycycline to induce recruitment of rTetR-effector fusion constructs to TetO-binding sites for 5 days (to test repression of reporter) or 2 days (to test activation of reporter). Fluorescence was analyzed by flow cytometry on a BD FACSCanto ll SORP.

### Luciferase Reporter Assays

HEK293T cells were co-transfected with a reporter plasmid (expressing Firefly luciferase under the control of a minCMV promoter with TetO sites), an effector plasmid (expressing an rTetR-effector fusion construct), and a normalization plasmid (expressing Renilla luciferase) by CaCl2-BBS transfection. Cells were also transfected with a plasmid expressing ZMYND11-WT, ZMYND11-R600W, or a BFP-only packaging plasmid. Transfected cells were treated with 1 μg/mL Doxycycline to induce recruitment of rTetR-effector fusion constructs to TetO-binding sites for 24 hours. Cells were washed with PBS and lysed with Passive Lysis Buffer (Promega). Firefly and Renilla luciferase activity were quantified using the Dual-Luciferase Reporter Assay (Promega).

### Small Molecule Drug Administration

Small molecule drugs (dTAG-13, dTAG^v^-1, revumenib, EZM-0414) were prepared fresh in DMSO to a 1000X concentration and stored at -20°C for up to three months. Primary cultured neurons were treated with a volume of 1000X drug stock equal to 1/1000 of culture volume. Defrosted stocks were not freeze-thawed. Control neurons were either treated with an equal concentration of inactive dTAG analogue (dTAG-13-NEG, dTAG^v^-1-NEG) or DMSO alone.

## Quantification and Analysis Methods

### RNA-Seq Quantification and Differential Expression

Reads were pseudoaligned to a transcriptome reference of the mm10 mouse reference genome using kallisto (v0.43.1) ^71^. Differential expression analysis was performed using DESeq2 to calculate significantly different transcript abundances between conditions ^72^. Significant genes were classified as those passing a log2 fold-change cutoff of ± 0.25 and an FDR-adjusted p-value of < 0.1.

### Pathway Enrichment Analysis

Lists of up- and down-regulated genes were generated using DESeq2 (described above). Pathway enrichment analysis of up- and down-regulated gene lists was performed using gProfiler2^73^. Queries were run with settings organism = "mmusculus", correction_method = "fdr".

### RNA Splicing Analysis

Transcript splicing analysis was performed using rMATS v4.1.1 using reads aligned to the mm10 mouse reference genome using STAR (v2.7.3a)^74,75^. Significantly enriched splicing events were quantified between *Zmynd11^fl/fl^;AAV-Cre* and *Zmynd11^fl/fl^;AAV-ΔCre* RNA libraries for the categories of Skipped Exons (SE), Intron Retention (IR), Mutually Exclusive Exons (MXE), Alternate 5’ Splice Site (A5SS), and Alternate 3’ Splice Site (A3SS). Only events called as significant on the basis of reads spanning exon-exon splice junctions were included.

### PRO-Seq Analysis

Random hexanucleotide barcodes contained within sequencing adaptors enabled removal of duplicate reads using BBTools Clumpify (v37.90)^76^. Reads were trimmed to 25 bp and mapped to the mm10 mouse reference genome using Bowtie2 (v2.2.9). Reads were converted to BED format using BEDTools (v2.26.0) and trimmed to the width of a single nucleotide at the second-to-3’ position, corresponding to the position of RNA Polymerase II prior to the incorporation of the terminal 11-biotin-NTP. PRO-Seq reads were mapped to the coordinates of transcription start sites (± 1000 bp) and gene bodies (minus TSS) using BEDOPS (v2.4.30). Gene annotations were sourced from GENCODE Release M25. For calculation of pausing index, only PRO-Seq reads mapping to the sense (coding) strand for a given gene were used.

### CUT&RUN Peak Calling

Peaks were called using MACS2 (v2.2.7.1) with parameters -f BAMPE -q 0.00001 --keep-dup all -g mm --call-summits^77^. Consensus ZMYND11 peaks were classified as those overlapping between two independent replicates of ZMYND11 CUT&RUN. To identify significantly enriched ZMYND11 peaks, ZMYND11 CUT&RUN signal was averaged across consensus peaks for two replicates of ZMYND11 CUT&RUN from wild-type cortical tissue (used for peak calling), and two replicates of ZMYND11 CUT&RUN from paired *Zmynd11-cKO* (*Zmynd11^fl/fl^;Baf53b-Cre*) cortical tissue, and the enrichment of signal in wild-type relative to cKO calculated. Consensus MLL1 peaks were classified as those overlapping between four independent replicates of MLL-N (KMT2A N-term) CUT&RUN, two independent replicates of RBBP5 CUT&RUN, and two independent replicates of ASH2L CUT&RUN.

### Peak Annotation and Motif Search

Genomic annotation of consensus ZMYND11 and MLL1 peaks was performed using HOMER (v4.9) annotatePeaks to categorize the genomic locus for each peak, the nearest gene, and to identify transcription factor motifs within peaks^78^. HOMER *de novo* motif results are reported separately for ZMYND11 peaks overlapping transcription start sites (TSS Peaks) and gene bodies (GB Peaks).

### Generation of Heatmap and Aggregate Plots

CUT&RUN signal was averaged across genomic loci using DeepTools (v3.0.2)^79^. To plot average signal across genes, DeepTools parameters computeMatrix scale-regions -- missingDataAsZero -m 4000 -a 2000 -b 2000 -bs 10. To plot average signal around peaks, DeepTools parameters computeMatrix reference-point --referencePoint center – missingDataAsZero -a 1000 -b 1000 --binSize 5. Aggregated data was plotted using the Python library matplotlib.

### Protein Mass Spectrometry Analysis

For spectral count mass spectrometry of endogenous FLAG-HA-ZMYND11 immunoprecipitates from brain tissue, unique peptide counts were combined from three independent replicates. Counts were compared between three paired replicates of *Zmynd11^FH/FH^* and *Zmynd11^WT/WT^* brain tissue. Proteins were plotted in a ranked scatter plot by the degree of enrichment in peptide counts for each immunoprecipitated protein in the *Zmynd11^FH/FH^* condition compared with wild-type controls.

For TMT-labeled multiplex mass spectrometry, two replicates were labeled and pooled per IP condition (FH-ZMYND11-WT, FH-ZMYND11-ΔMYND, FH-ZMYND11-R600W, FH-GFP-SV40NLS). Intensities of identified proteins were measured and normalized relative to the intensity of bait protein for each immunoprecipitation experiment. FH-ZMYND11-WT protein intensities were quantified relative to GFP control, and mutant ZMYND11 protein intensities (ΔMYND and R600W) were quantified relative to ZMYND11-WT. Proteins with fewer than two peptides quantified were excluded from analysis.

### Flow Cytometry Analysis

Flow cytometry data was quantified using Cytoflow (v1.2) and custom Python scripts^80^. Gating was applied to select events representing viable single cells expressing mCherry. Thresholds of Citrine fluorescence were set on the basis of untreated rTetR-only expressing cells.

### Reagents and Resources

**Antibodies**
|  |  |  |  |
| --- | --- | --- | --- |
| 1323 | Guinea Pig anti-Rabbit | Fisher | Cat# NBP172763 |
| 1324 | Rabbit anti-Histone H3K4me1 | Abcam | Cat# ab176877 |
| 1325 | Rabbit anti-Histone H3K4me3 | Abcam | Cat# ab213224 |
| 1326 | Rabbit anti-Histone H3K27ac | Abcam | Cat# ab4729 |
| 1327 | Rabbit anti-Histone H3K27me3 | Abcam | Cat# ab192985 |
| 1328 | Rabbit anti-Histone H3K36me3 | Abcam | Cat# ab9050 |
| 1329 | Rabbit anti-Histone H3 | Abcam | Cat# ab1791 |
| 1330 | Rabbit anti-ASH2L | Abcam | Cat# ab176334 |
| 1331 | Rabbit anti-KMT2A/MLL1 (C-term) | CST | Cat# 14197 |
| 1332 | Rabbit anti-KMT2A/MLL1 (N-term) | CST | Cat# 14689 |
| 1333 | Rabbit anti-KMT2B/MLL2 (N-term) | CST | Cat# 63735 |
| 1334 | Rabbit anti-MENIN | CST | Cat# 19893 |
| 1335 | Rabbit anti-RBBP5 | Active Motif | Cat# 61406 |
| 1336 | Rabbit anti-TBP | CST | Cat# 44059 |
| 1337 | Rabbit anti-WDR5 | CST | Cat# 13105 |
| 1338 | Rabbit anti-ZMYND11 | Abcam | Cat# ab190890 |
| 1339 | Rabbit anti-ZMYND11 | CST | Cat# 67713 |
| 1340 | Rabbit anti-HA | CST | Cat# 3724 |
| 1341 | Rat anti-HA | Roche | Cat# 11867423001 |
| 1342 | Normal Rabbit IgG | CST | Cat# 2729 |
| 1343 |  |  |  |
| 1344 | **Commercial Assays** |  |  |
| 1345 | Dual-Luciferase Reporter Assay System | Promega | Cat# E1980 |
| 1346 | FD Rapid GolgiStain | FD Neurotech | Cat# PK401 |
| 1347 | NEBNext Ultra Directional Library Prep Kit | NEB | Cat# E7420L |
| 1348 | NEBNext rRNA Depletion Kit | NEB | Cat# E6310X |
| 1349 | NEBNext Multiplex Oligos for Illumina | NEB | Cat# E7335L |
| 1350 | RNeasy Micro Kit | QIAGEN | Cat# 74004 |
| 1351 | ThruPLEX DNA-Seq Kit | Takara | Cat# R400676 |
| 1352 |  |  |  |
| 1353 | **Enzymes** |  |  |
| 1354 | Micrococcal Nuclease | NEB | Cat# M0247 |
| 1355 | RNA 5 Pyrophosphohydrolase (RppH) | NEB | Cat# M0356 |
| 1356 | Q5 High-Fidelity DNA Polymerase | NEB | Cat# M0491 |
| 1357 | Superscript III Reverse Transcriptase | Invitrogen | Cat# 18080044 |
| 1358 | T4 Polynucleotide Kinase | NEB | Cat# M0201 |
| 1359 | T4 RNA Ligase 1 | NEB | Cat# M0204 |
| 1360 |  |  |  |
| 1361 | **Other Reagents** |  |  |
| 1362 | BioMag Plus Concanavalin A Beads | Bangs | Cat# BP531 |
| 1363 | Eukitt Quick-Hardening Mounting Medium | Sigma Aldrich | Cat# 03989 |
| 1364 | GlycoBlue Coprecipitant | Invitrogen | Cat# AM9515 |
| 1365 | Hydrophilic Streptavidin Magnetic Beads | NEB | Cat# S1421 |
| 1366 | Pierce Anti-DYKDDDDK Magnetic Agarose | Life Tech | Cat# A36797 |
| 1367 | Pierce Protein A/G Magnetic Agarose Beads | Life Tech | Cat# 78609 |
| 1368 | SUPERase-In RNase Inhibitor | Invitrogen | Cat# AM2694 |
| 1369 | SuperSignal West Pico PLUS | Thermo | Cat# 34580 |
| 1370 |  |  |  |
| 1371 | **Experimental Model Strains** |  |  |
| 1372 | Mouse: *Zmynd11-fl* | This Study |  |
| 1373 | Mouse: *Zmynd11-FLAG-HA* | This Study |  |
| 1374 | Mouse: *Zmynd11-dTAG* | This Study |  |
| 1375 | Mouse: Baf53b-Cre | The Jackson Laboratory | Strain: 027826 |
| 1376 | Mouse: E2a-Cre | The Jackson Laboratory | Strain: 003724 |

