## Supplementary material for "ZMYND11 Restrains KMT2A to Enable a Neuronal Developmental Program": supp_figures_with_legends

Supplementary Figure 1

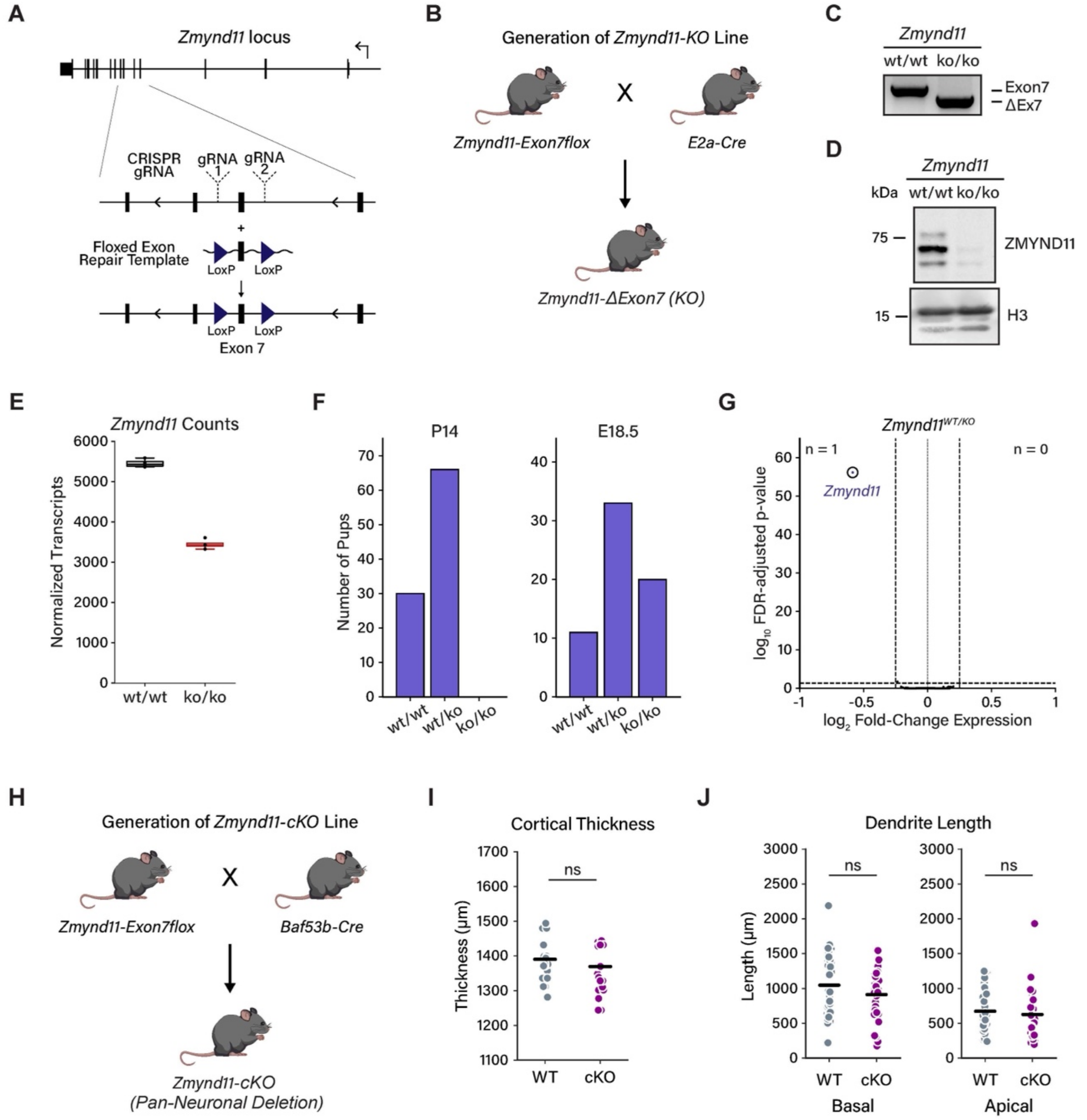


**Supplementary Figure 1. Germline disruption of *ZMYND11* impacts mouse development in homozygotes but not heterozygotes**

a) Schematic showing generation of *Zmynd11-Exon7Flox* (*Zmynd11$^{fl}$*) mouse line using CRISPR-Cas9 and homologous recombination with a single-stranded DNA repair template.

b) Schematic showing generation of constitutively recombined *Zmynd11-ΔExon7* (*Zmynd11-KO*) mice by crossing *Zmynd11$^{fl}$* to the germline-expressing *E2a-Cre*.

c) PCR genotyping showing Cre-mediated excision of *Zmynd11* exon 7 in the germline in descendants of *E2a-Cre*-expressing *Zmynd11$^{fl}$* mice.

d) Western blot showing ZMYND11 protein in wild-type and *Zmynd11*$^{KO/KO}$ E18.5 cortex, with Histone H3 as loading control.
e) Box plot of DESeq2-normalized transcript counts of *Zmynd11* in RNA libraries generated from the cortex of 6-week-old *Zmynd11*$^{+/-}$ and wild-type mice; n = 4 individual mice per condition.
f) Distribution of genotype frequencies in *Zmynd11-KO* litters that were tail-clipped at P14 (left panel) or dissected at E18.5 (right panel).
g) Volcano plot showing DESeq2-identified gene expression changes in whole cortex from 6-week-old *Zmynd11*$^{+/-}$ vs wild-type mice. Plot shows $\log_2$ fold-change RNA expression vs. $-\log_{10}$ p-value with Benjamini-Hochberg false discovery rate (FDR) adjustment. n = 4 individual mice per condition.
h) Schematic showing generation of neuron-specific conditional knockout *Zmynd11-fl;Baf53b-Cre* (*Zmynd11-cKO*) mice by crossing *Zmynd11-fl* to the neuronal-expressing *Baf53b-Cre* line.
i) Comparison of cortical thickness in 6-week-old *Zmynd11*$^{fl/fl}$*; Baf53b-Cre* and *Zmynd11*$^{wt/wt}$*; Baf53b-Cre* mice. ns = not significant, p-values calculated using Wilcoxon rank-sum test; n = 6 individual mice per condition.
j) Comparison of dendrite length in 6-week-old *Zmynd11*$^{fl/fl}$*; Baf53b-Cre* and *Zmynd11*$^{wt/wt}$*; Baf53b-Cre* mice, separated by apical and basal dendrites. ns = not significant, p-values calculated using Wilcoxon rank-sum test; n = 6 cells per mouse, 6 individual mice per condition.

Supplementary Figure 2

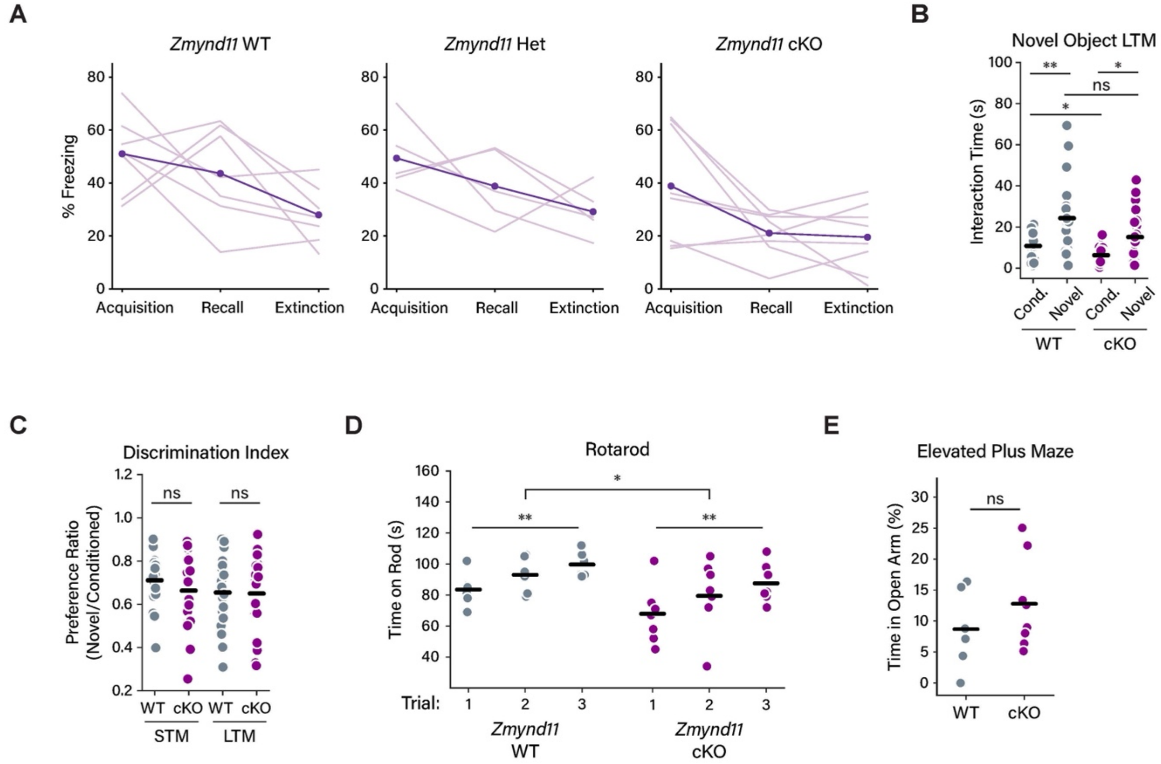


**Supplementary Figure 2. Behavioral testing of *Zmynd11-cKO* mice**

a) Quantification of freezing behavior during contextual fear conditioning, plotted separately by genotype showing the responses of individual mice.

b) Duration of mouse interactions with a novel and habituated object present in the same housing context, long-term recall. * = $p < 0.01$, ** = $p < 0.001$, p-values calculated using Wilcoxon rank-sum test; n = 19 individual mice (WT), n = 21 individual mice (cKO).

c) Ratio of preference between novel and conditioned objects in short- and long-term memory tasks for novel object recognition. ns = not significant. p-values calculated using Wilcoxon rank-sum test; n = 19 individual mice (WT), n = 21 individual mice (cKO).

d) Duration of mice balancing on rotating rod in Rotarod task. * = $p < 0.05$, ** = $p < 0.01$; p-values calculated using repeated measures ANOVA; n = 6 individual mice (WT), n = 9 individual mice (cKO).

e) Relative time spent by mice in open arm of elevated plus maze. ns = not significant, p-values calculated using Wilcoxon rank-sum test; n = 6 individual mice (WT), n = 9 individual mice (cKO).

Supplementary Figure 3

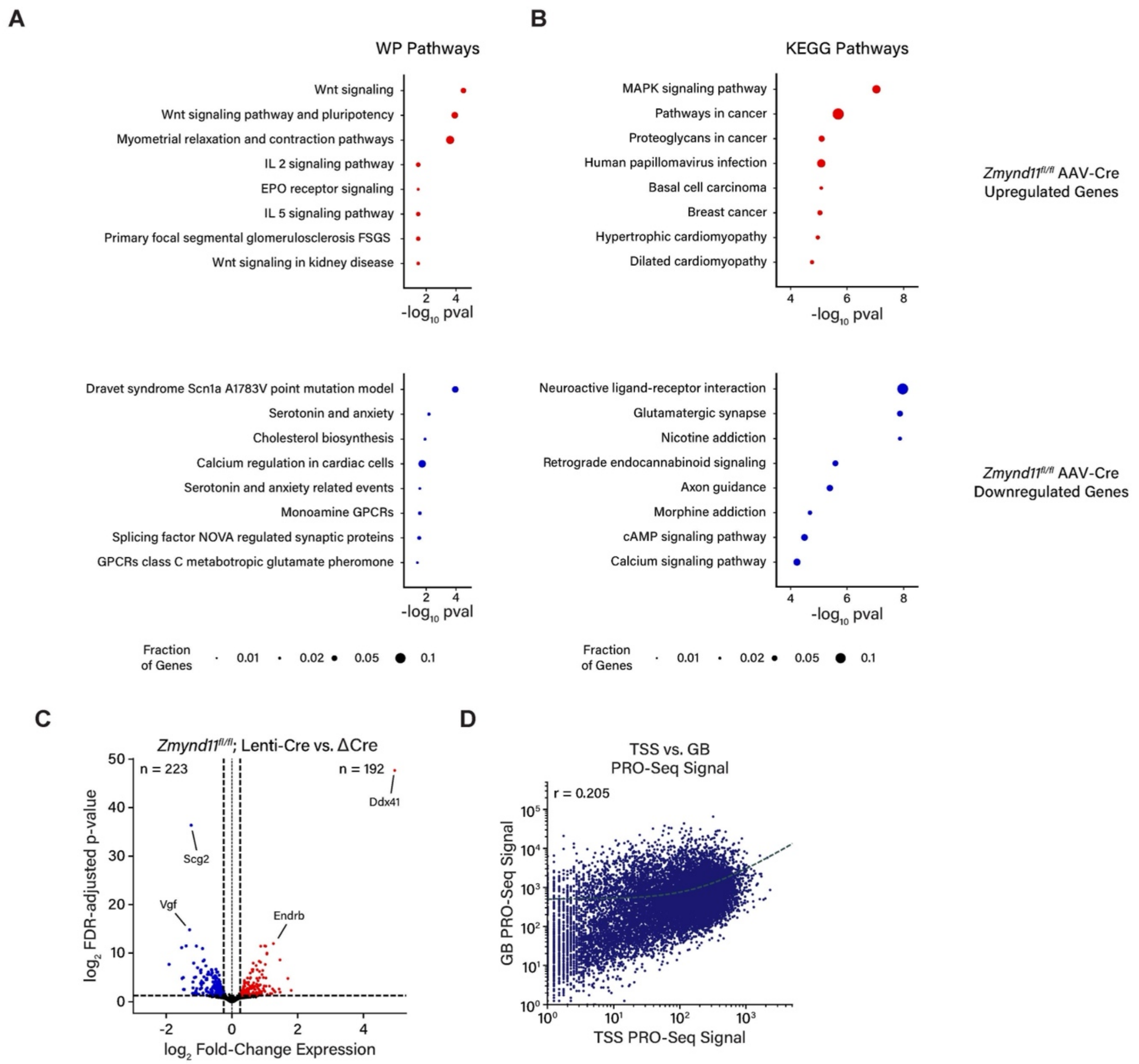


**Supplementary Figure 3. Analysis of gene expression in *Zmynd11-cKO***

a) Gene ontology analysis of DESeq2-identified significantly up- and down-regulated genes in *Zmynd11*$^{fl/fl}$ AAV-Cre vs. AAV-ΔCre cortex. WP (Wikipathways) gene set, analysis by gProfiler2.

b) Gene ontology analysis of DESeq2-identified significantly up- and down-regulated genes in *Zmynd11*$^{fl/fl}$ AAV-Cre vs. AAV-ΔCre cortex. KEGG Pathways gene set, analysis by gProfiler2.

c) Volcano plot of DESeq2 gene expression changes in *Zmynd11*$^{fl/fl}$ *Lenti-Cre* vs. *Lenti-ΔCre* DIV7 cortical cultures, plotting $\log_2$ fold-change RNA expression vs. $-\log_{10}$ p-value with Benjamini-Hochberg false discovery rate (FDR) adjustment (n = 3 individual dissections).

d) Scatter plot showing correlation of PRO-Seq read density at the transcription start site (TSS) vs. gene body (GB) for expressed genes in *Zmynd11*$^{fl/fl}$; *Lenti-ΔCre* DIV7 cortical cultures. r = Pearson correlation coefficient.

Supplementary Figure 4

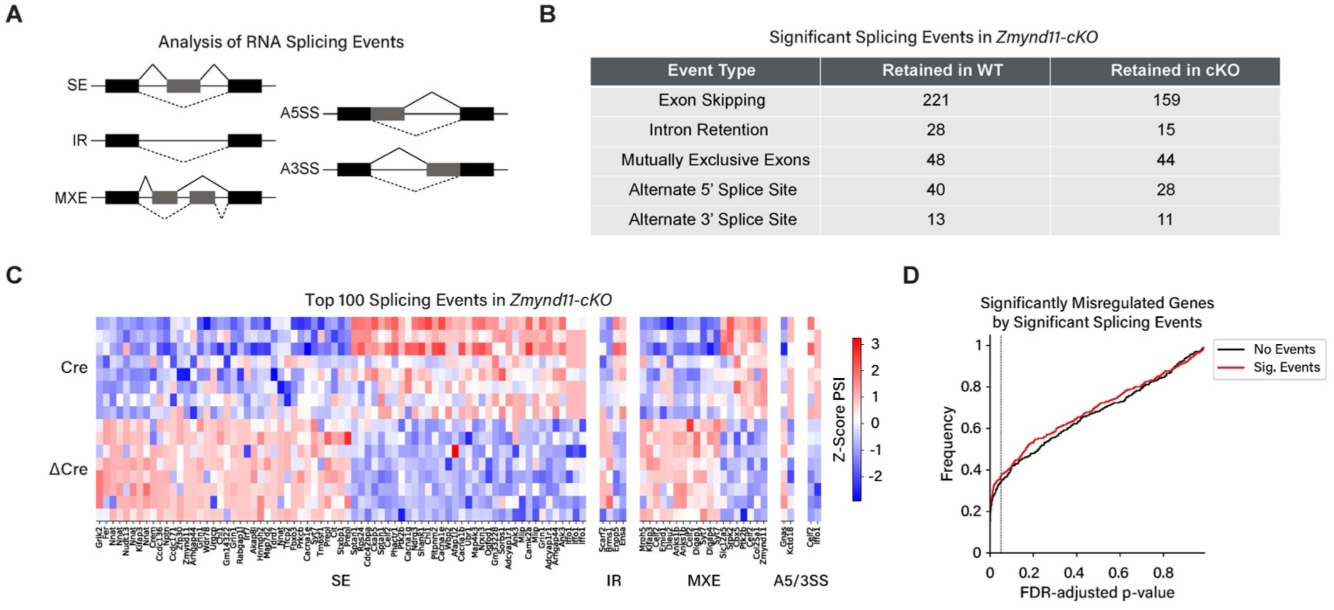


| Event Type | Retained in WT | Retained in cKO |
|---|---|---|
| Exon Skipping | 221 | 159 |
| Intron Retention | 28 | 15 |
| Mutually Exclusive Exons | 48 | 44 |
| Alternate 5' Splice Site | 40 | 28 |
| Alternate 3' Splice Site | 13 | 11 |

**Supplementary Figure 4. Analysis of alternative splicing in *Zmynd11-cKO***

a) Schematic showing five types of alternative splicing events quantified by Multivariate Analysis of Transcript Splicing with Replicates (rMATS). Skipped Exon (SE), Intron Retention (IR), Mutually Exclusive Exons (MXE), Alternative 5' Splice Site (A5SS), Alternative 3' Splice Site (A3SS).

b) Table of differential splicing events in *Zmynd11*$^{fl/fl}$ AAV-Cre vs. AAV-ΔCre cortex by rMATS. p-values calculated using likelihood-ratio test, n = 8 individual mice.

c) Heatmap of Z-Score Percent Spliced In (PSI) for top 100 significantly regulated alternative splicing events in *Zmynd11*$^{fl/fl}$ AAV-Cre vs. AAV-ΔCre cortex.

d) Cumulative histogram showing DESeq2 FDR-adjusted p-values for differential gene expression in *Zmynd11*$^{fl/fl}$ AAV-Cre vs. AAV-ΔCre, separated by genes with (red line) and without (black line) significant splicing events. Dotted line indicates significance cutoff of padj < 0.05.

Supplementary Figure 5

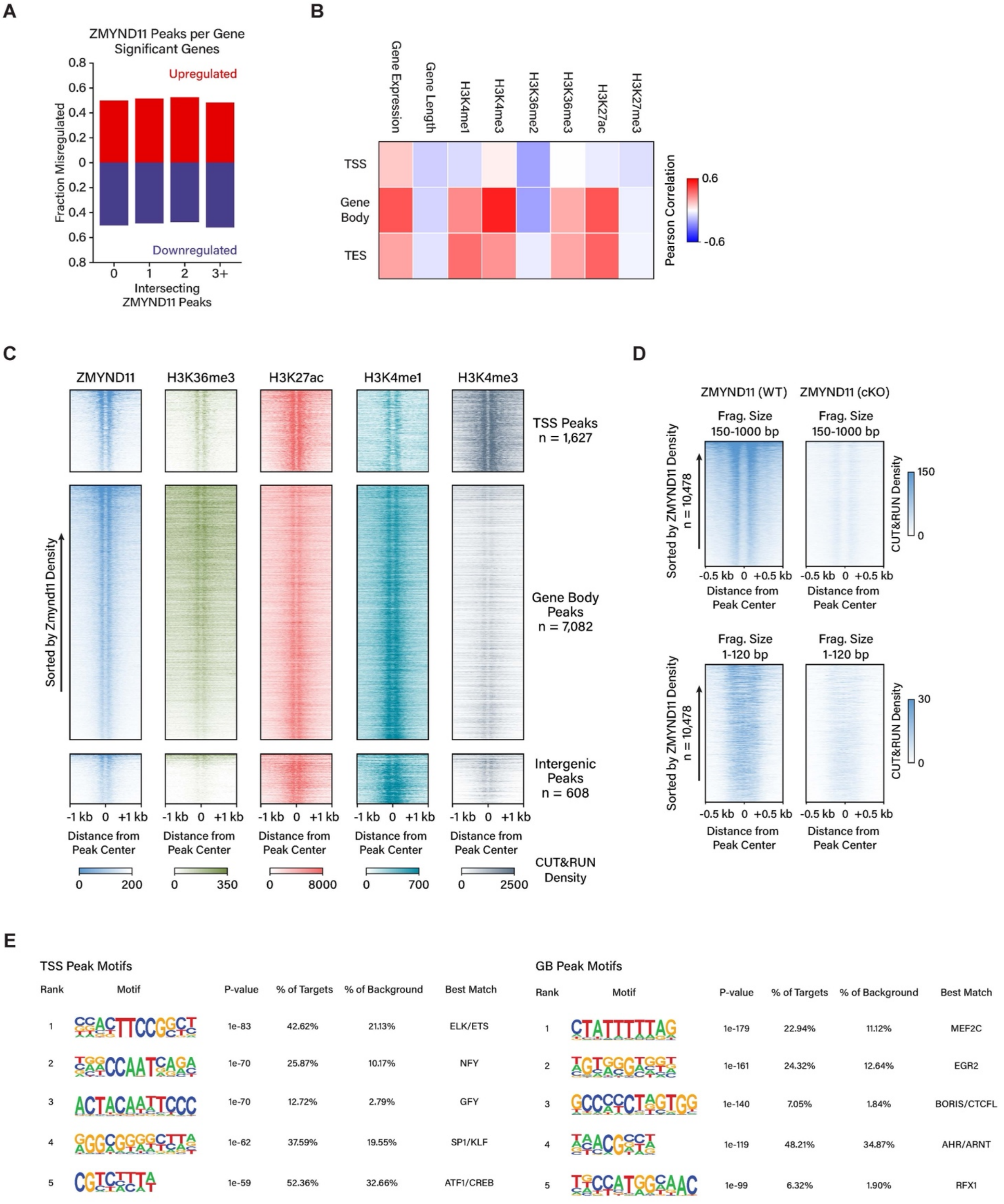

**Supplementary Figure 5. Analysis of ZMYND11 genomic binding patterns**

a) Proportion of genes which are significantly up- or down-regulated in *Zmynd11*$^{fl/fl}$ AAV-Cre vs. AAV-ΔCre cortex, grouped by the number of significant ZMYND11 CUT&RUN peaks per gene.

b) Heatmap showing Pearson correlation coefficient of ZMYND11 CUT&RUN signal with genomic features such as gene expression (transcripts per million), gene length, and CUT&RUN signal for histone modifications. Correlations are plotted for transcription start sites (TSS), gene body, and transcription end sites (TES).

c) Heatmaps of CUT&RUN signal (scaled read coverage per bp) for select histone marks surrounding ZMYND11 peaks identified by MACS2 at TSS, gene bodies, and intergenic regions.

d) Heatmaps of CUT&RUN signal density (scaled read coverage per bp) at ZMYND11 peaks, separated by fragment length into nucleosomal (150-1000 bp, top) and sub-nucleosomal (1-120 bp, bottom) fractions. CUT&RUN density of ZMYND11 shown for 6-week-old wild-type and ZMYND11-cKO mouse cortex; n = 2 individual mice per condition.

e) Motif analysis of DNA sequences within ZMYND11 peaks identified by MACS2 at transcription start sites (left) and gene bodies (right). De novo motif analysis by HOMER v4.9.

Supplementary Figure 6

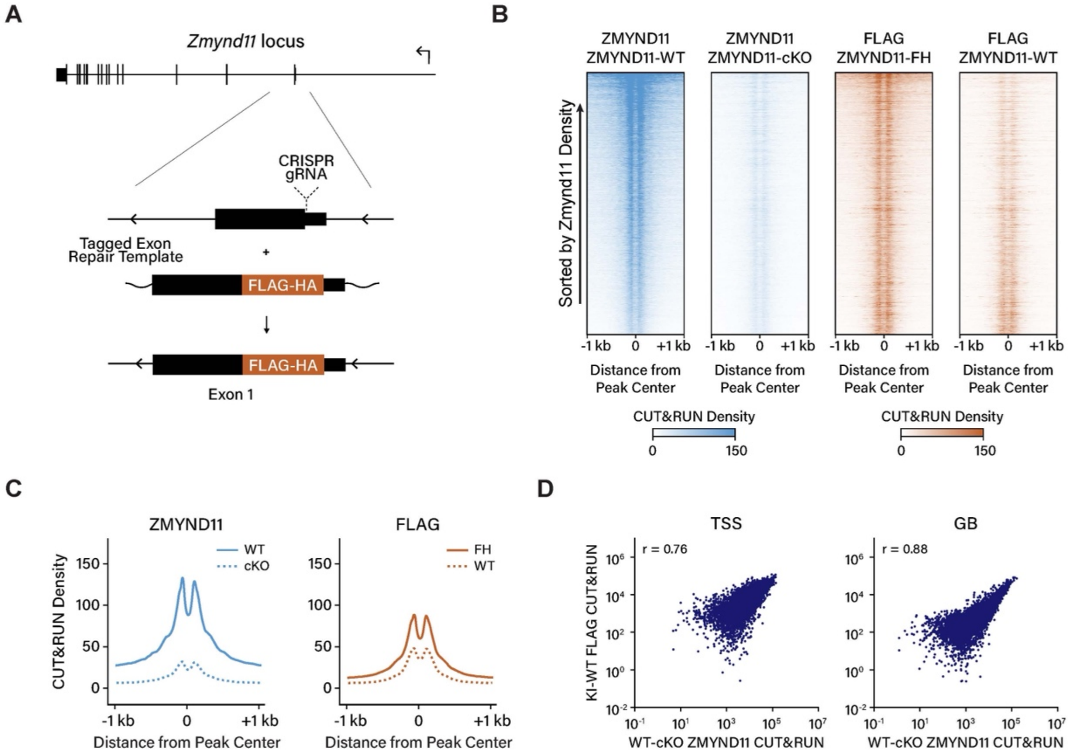


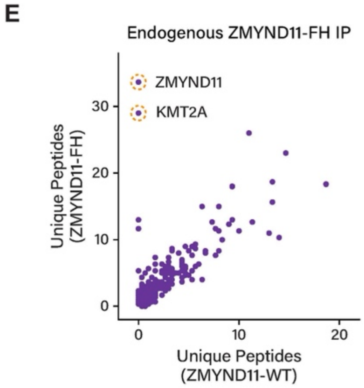


F

| Protein ID | Unique Peptides WT Rep1 | Unique Peptides FH Rep1 | Unique Peptides WT Rep2 | Unique Peptides FH Rep2 | Unique Peptides WT Rep3 | Unique Peptides FH Rep3 |
|---|---|---|---|---|---|---|
| ZMYND11 | 0 | 26 | 0 | 42 | 0 | 33 |
| KMT2A | 0 | 14 | 0 | 54 | 0 | 19 |
| ZBTB38 | 0 | 7 | 0 | 26 | 0 | 6 |
| NSD1 | 0 | 2 | 0 | 16 | 0 | 17 |
| BCORL1 | 0 | 0 | 0 | 11 | 0 | 5 |
| NUMA1 | 0 | 2 | 0 | 0 | 1 | 11 |
| POLR2A | 0 | 2 | 0 | 0 | 2 | 13 |
| CHD5 | 0 | 1 | 0 | 1 | 3 | 15 |
| TFIP11 | 0 | 1 | 0 | 0 | 5 | 21 |
| POLR2B | 0 | 1 | 0 | 0 | 1 | 9 |
| PSIP1 | 0 | 2 | 2 | 10 | 3 | 7 |
| HNRNPR | 0 | 0 | 0 | 0 | 0 | 7 |
| CSNK2A1 | 0 | 0 | 0 | 0 | 0 | 7 |
| CRNKL1 | 0 | 0 | 0 | 0 | 1 | 9 |

**Supplementary Figure 6. Generation and validation of *ZMYND11-FH* mouse line**

a) Schematic showing generation of the epitope-tagged Zmynd11 (*Zmynd11-FH*) mouse line using CRISPR-Cas9 and homologous recombination with a single-stranded DNA repair template.

b) Heatmaps of ZMYND11 and FLAG CUT&RUN signal density (scaled read coverage per bp) at ZMYND11 peaks from 6-week-old mouse cortex. ZMYND11 CUT&RUN density shown for wild-type and control *Zmynd11-cKO* samples; FLAG CUT&RUN density shown for *Zmynd11-FH* and control wild-type samples. n = 2 individual mice per condition.

c) Aggregate plots showing average distribution of ZMYND11 and FLAG CUT&RUN signal centered around significant ZMYND11 peaks (MACS no-control p-value < 1e-05, ZMYND11 CUT&RUN WT/cKO > 2), summed signal from (b).

d) Scatter plot showing correlation of CUT&RUN signal intensity for ZMYND11 (in wild-type, normalized vs. *Zmynd11-cKO*) and FLAG (in *Zmynd11-FH*, normalized vs. wild-type) at transcription start sites (left) and gene bodies (right). r = Pearson correlation coefficient; n = 2 individual mice per condition.

e) Scatter plot showing proteins identified by mass spectrometry in eluates of anti-FLAG immunoprecipitation experiments using cortical lysates from *Zmynd11-FH* and wild-type mice, n = 3 independent replicates per condition.

f) Spectral counts of proteins identified by mass spectrometry in eluates of anti-FLAG immunoprecipitation experiments using cortical lysates from *Zmynd11-FH* and wild-type mice; n = 3 independent replicates per condition.

Supplementary Figure 7

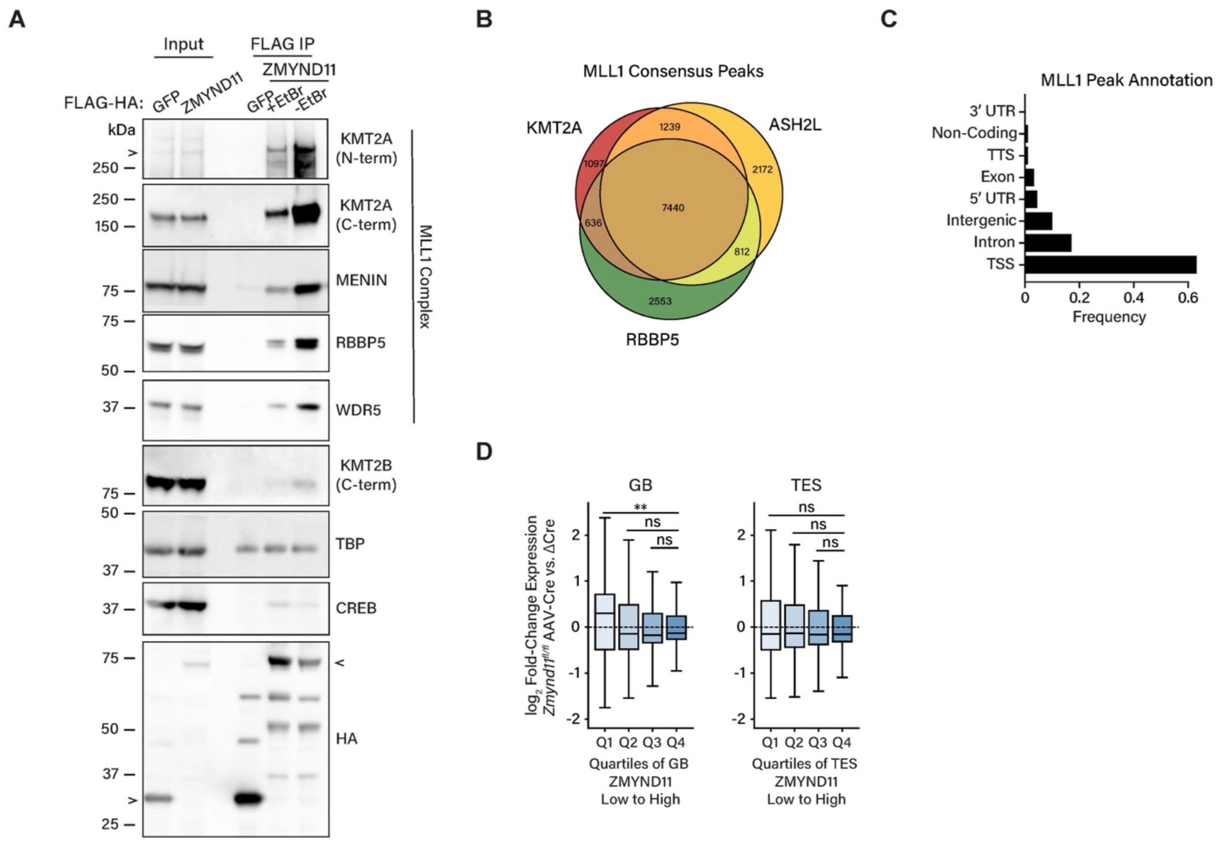


**Supplementary Figure 7. Association of ZMYND11 with KMT2A**

a) Representative western blot showing co-immunoprecipitation of ZMYND11, KMT2A, and MLL complex members from nuclear lysates of HEK293T cells expressing FLAG-tagged ZMYND11 or GFP, with and without ethidium bromide (EtBr) treatment. Data are representative of three independent experiments.

b) Venn diagram showing overlap of CUT&RUN peaks (MACS no-control P-value < 1E-05) for KMT2A, RBBP5, ASH2L to define MLL1 consensus peaks.

c) Annotation of the genomic localization of consensus MLL1 peaks defined in (b) by overlap of KMT2A, RBBP5, and ASH2L (Homer annotatePeaks).

d) Box plots showing differential gene expression by DESeq2 analysis of RNA-seq in *Zmynd11*$^{fl/fl}$ AAV-Cre vs. AAV-ΔCre, with genes separated into quartiles based on enrichment of WT/cKO ZMYND11 CUT&RUN signal at the gene body (left) and TES (right) (low to high). ns = not significant, ** = $p < 0.01$, p-values calculated using Wilcoxon rank-sum test; n = 3 independent replicates per condition.

Supplementary Figure 8

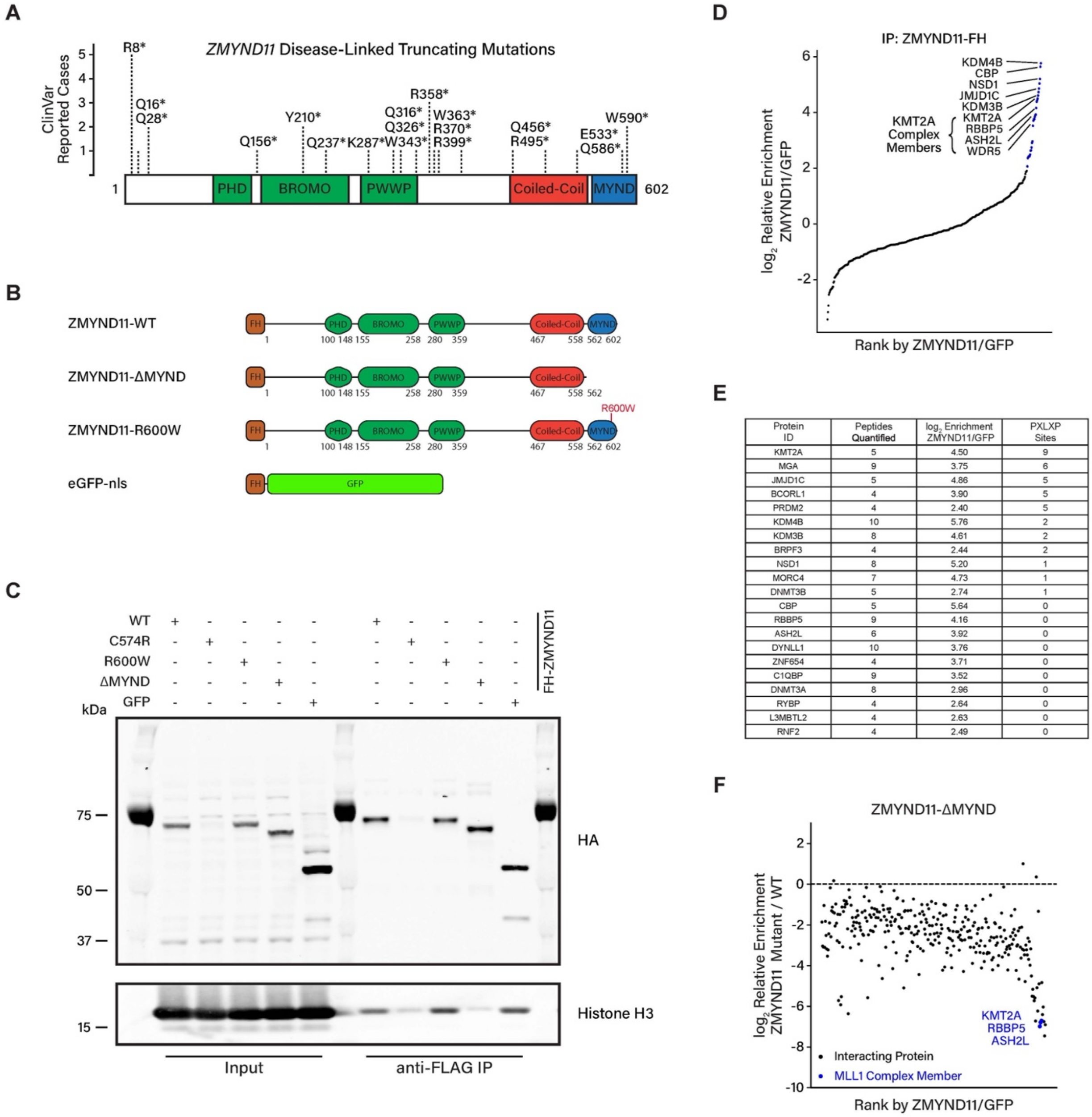


| Protein ID | Peptides Quantified | $\log_2$ Enrichment ZMYND11/GFP | PXLXP Sites |
|---|---|---|---|
| KMT2A | 5 | 4.50 | 9 |
| MGA | 9 | 3.75 | 6 |
| JMJD1C | 5 | 4.86 | 5 |
| BCORL1 | 4 | 3.90 | 5 |
| PRDM2 | 4 | 2.40 | 5 |
| KDM4B | 10 | 5.76 | 2 |
| KDM3B | 8 | 4.61 | 2 |
| BRPF3 | 4 | 2.44 | 2 |
| NSD1 | 8 | 5.20 | 1 |
| MORC4 | 7 | 4.73 | 1 |
| DNMT3B | 5 | 2.74 | 1 |
| CBP | 5 | 5.64 | 0 |
| RBBP5 | 9 | 4.16 | 0 |
| ASH2L | 6 | 3.92 | 0 |
| DYNLL1 | 10 | 3.76 | 0 |
| ZNF654 | 4 | 3.71 | 0 |
| C1QBP | 9 | 3.52 | 0 |
| DNMT3A | 8 | 2.96 | 0 |
| RYBP | 4 | 2.64 | 0 |
| L3MBTL2 | 4 | 2.63 | 0 |
| RNF2 | 4 | 2.49 | 0 |

**Supplementary Figure 8. Screening interactions of ZMYND11 variants associated with neurodevelopmental disorders**

a) Schematic showing truncating mutations found in cases of *ZMYND11*-linked neurodevelopmental disorder in human patients, from ClinVar reports evaluated ‘Pathogenic’ and ‘Likely Pathogenic’.

b) Schematic showing the domain architecture of the ZMYND11 variants and GFP control used in HEK293T co-immunoprecipitation experiments in Fig. 5b, Fig S8c-f.

c) Representative western blot showing anti-FLAG immunoprecipitation of FLAG-HA-tagged ZMYND11 constructs expressed in HEK293T cells, blotting back with an anti-HA antibody, with Histone H3 as a loading control. The ZMYND11 variant C574R is not stable when over-expressed. Data are representative of two independent experiments.

d) Ranked plot of protein hits by relative enrichment in ZMYND11-FH IP compared to GFP-FH IP in TMT-IP-MS from HEK 293T cells. MLL1 complex members are indicated in blue; n = 2 independent replicates per condition.

e) List of interacting proteins showing >4-fold enrichment in Zmynd11-FH IP over GFP-FH IP, along with the number of peptides identified in the TMT-MS experiment and the number of PXLXP motifs present within the full corresponding protein sequence.

f) Scatter plot showing the relative enrichment of co-immunoprecipitating proteins with FLAG-tagged ZMYND11-ΔMYND compared to wild-type ZMYND11, ordered by relative enrichment in wild-type vs. FLAG-tagged GFP control. MLL1 Complex members are indicated in blue; n = 3 independent transfections per condition.

Supplementary Figure 9

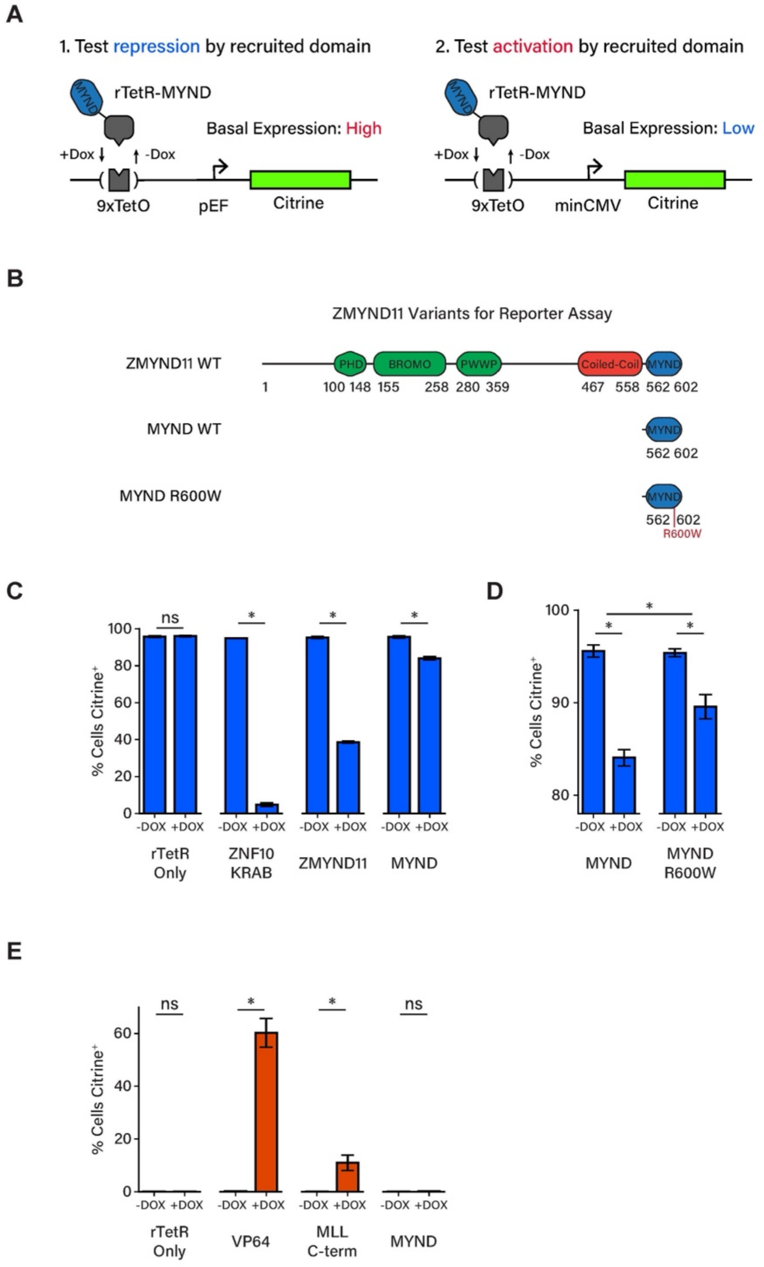


**Supplementary Figure 9. Repression of reporter gene expression by ZMYND11**

a) Schematic showing implementation of fluorescent reporter assay for repression or activation of reporter gene by recruitment of rTetR-fusion constructs with the ZMYND11 MYND domain or other effectors.

b) Schematic showing the domain architecture of ZMYND11 variants assayed for effects on reporter expression.

c) Bar plots showing reporter assay for repression. Values reflect the percentage of Citrine-positive cells after treatment with DMSO (-DOX) or 1 µg/mL doxycycline (+DOX) for 5 days. Constructs: Control (rTetR alone), ZNF10-KRAB (positive control for repression), ZMYND11, ZMYND11 MYND domain. Mean values are shown ± SD; n = 2 independent transductions per condition. ns = not significant, ** = $p < 0.05$, p-values calculated using Wilcoxon rank-sum test.

d) Bar plots showing reporter assay for repression. Values reflect the percentage of Citrine-positive cells after treatment with DMSO (-DOX) or 1 µg/mL doxycycline (+DOX) for 5 days. Constructs: ZMYND11 MYND domain (wild-type), ZMYND11 MYND domain (R600W). Mean values are shown ± SD; n = 2 independent transductions per condition. ns = not significant, ** = $p < 0.05$, p-values calculated using Wilcoxon rank-sum test.

e) Bar plots showing reporter assay for activation. Values reflect the percentage of Citrine-positive cells after treatment with DMSO (-DOX) or 1 µg/mL doxycycline (+DOX) for 5 days. Constructs: Control (rTetR alone), VP64 (positive control for activation), MLL-C (KMT2A C-terminal region), ZMYND11 MYND domain. Mean values are shown ± SD; n = 2 independent transductions per condition. ns = not significant, ** = $p < 0.05$, p-values calculated using Wilcoxon rank-sum test.

Supplementary Figure 10

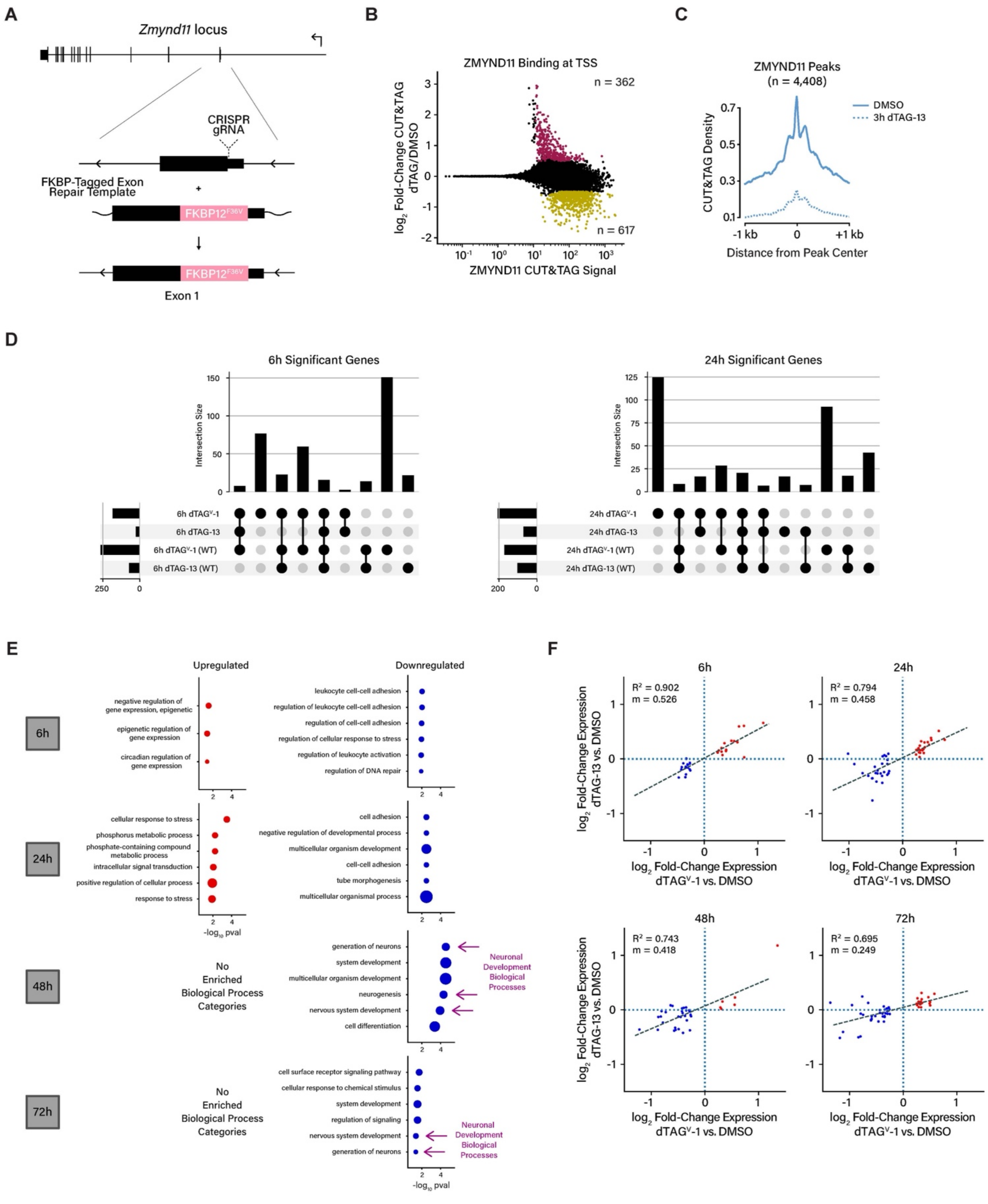

**Supplementary Figure 10. Generation and characterization of *Zmynd11-dTAG* mouse line**

a) Schematic showing the generation of the $FKBP12^{F36V}$-tagged (*Zmynd11-dTAG*) mouse line using CRISPR-Cas9 and homologous recombination with a single-stranded DNA template.

b) Scatter plot showing ZMYND11 CUT&TAG signal at transcription start sites in *Zmynd11-dTAG* cultured primary cortical neurons treated with 50 nM dTAG-13 for 3 hours, relative to DMSO vehicle control. Genes showing significant increase or decrease in signal are highlighted (significance calculations by DESeq2, FDR-adjusted pval < 0.1, minimum $\log_2$ fold-change 0.5); n = 5 independent replicates.

c) Aggregate plot showing average distribution of ZMYND11 CUT&TAG signal centered around significant ZMYND11 peaks, $Zmynd11^{dTAG/dTAG}$ cultured primary cortical neurons treated with 500 nM dTAG-13 or DMSO vehicle for 3 hours (MACS2 no-control p-value < 1e-05, ZMYND11 CUT&RUN DMSO/dTAG > 2).

d) Intersection of genes exhibiting significant expression differences in $Zmynd11^{dTAG/dTAG}$ or $Zmynd11^{WT/WT}$ cultured primary cortical neurons treated with 500 nM dTAG-13 or $dTAG^{v}$-1 for 6 or 24 hours, relative to DMSO control. Significance calculations by DESeq2, FDR-adjusted pval < 0.1, minimum $\log_2$ fold-change 0.5, n = 4 independent replicates for $Zmynd11^{dTAG/dTAG}$ neurons, n = 2 independent replicates for $Zmynd11^{WT/WT}$ neurons.

e) Gene ontology analysis of DESeq2 significantly up- and down-regulated genes in $Zmynd11^{dTAG/dTAG}$ neurons treated with $dTAG^{v}$-1, GO:BP (Biological Process) gene set, analysis by gProfiler2.

f) Scatter plot showing relative differential expression in $Zmynd11^{dTAG/dTAG}$ cultured primary cortical neurons treated for 24 hours with 500 nM dTAG-13 or $dTAG^{v}$-1 vs. DMSO control, gene list selected on the basis of significant differential expression in $dTAG^{v}$-1 vs. DMSO condition.

### Supplementary Figure 11

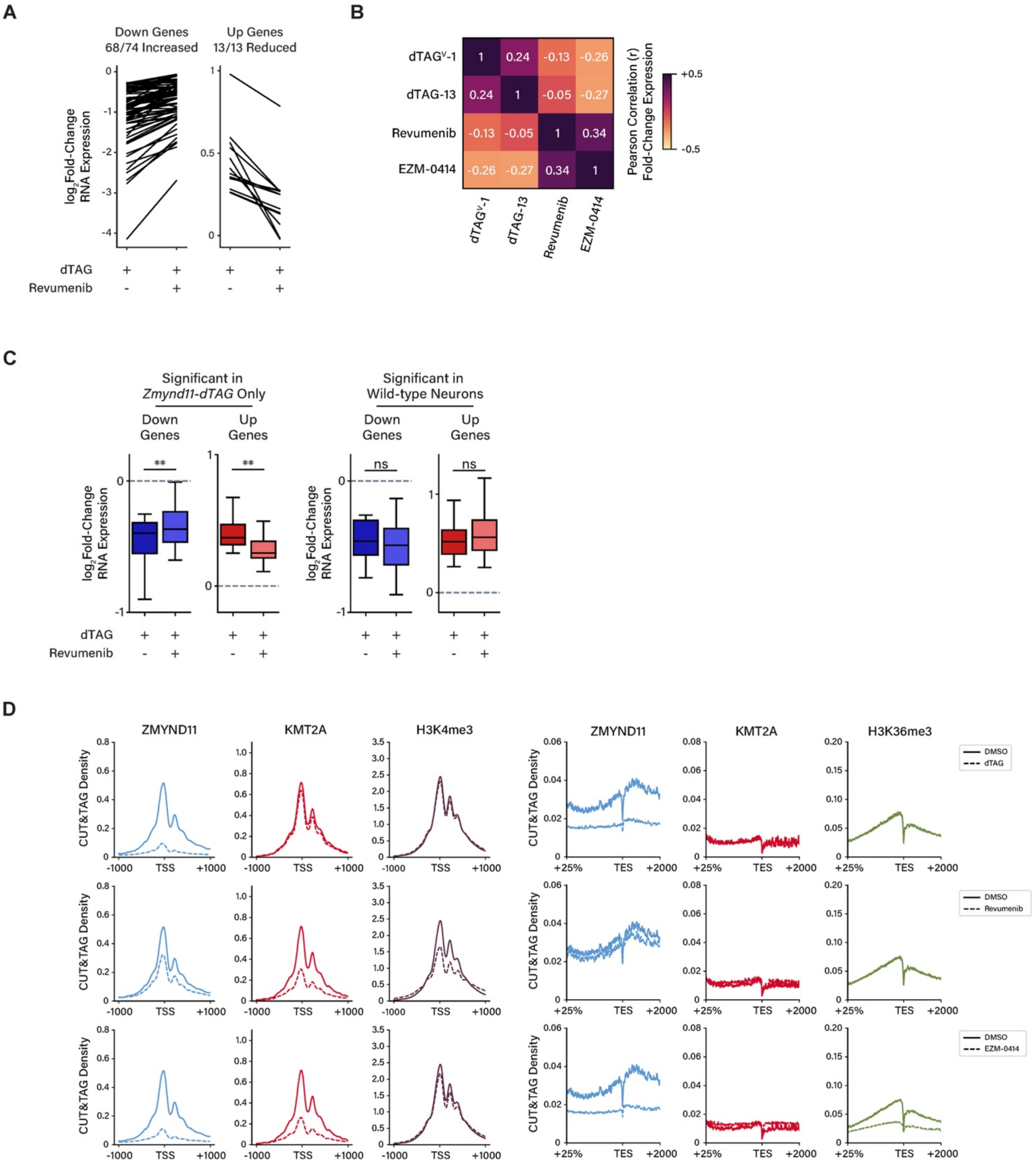

**Supplementary Figure 11. Interrogation of ZMYND11 function and binding using inhibitors of KMT2A and SETD2.**

a) Line plot showing the change in expression of each individual gene significantly misregulated in *Zmynd11-dTAG* cultured primary cortical neurons with 500 nM dTAG$^{v}$-1 administration for 24 hours, showing fold-change in expression upon dTAG treatment with or without 2 μM revumenib relative to 500 nM dTAG$^{v}$-1-NEG control. Each line represents an individual gene.

b) Heatmap showing Pearson correlation (r) of fold-change gene expression in *Zmynd11-dTAG* cultured primary cortical neurons treated with 500 nM dTAG$^{v}$-1, 500 nM dTAG-13, 2 μM revumenib, and 2 μM EZM-0414 for 24 hours. dTAG-13 and dTAG$^{v}$-1 relative to dTAG-13-NEG and dTAG$^{v}$-1-NEG control, respectively, Revumenib and EZM-0414 relative to DMSO control. Fold-change values calculated from four independent RNA-seq replicates.

c) Change in expression of genes significantly misregulated in *Zmynd11-dTAG* primary cortical neurons with 500 nM dTAG$^{v}$-1 administration for 24 hours relative to DMSO control, with and without co-administration of 2 μM revumenib. Genes separated by those significantly changed only in *Zmynd11$^{dTAG/dTAG}$* neurons, and those significantly changed in both *Zmynd11$^{dTAG/dTAG}$* (left) and *Zmynd11$^{WT/WT}$* neurons (right). ns = not significant, ** = $p < 0.001$, p-values calculated using Wilcoxon rank-sum test; n = 4 independent RNA-seq replicates (*Zmynd11$^{dTAG/dTAG}$* neurons), n = 2 independent RNA-seq replicates (WT neurons).

d) Aggregate plots of ZMYND11, KMT2A, and H3K36me3 CUT&TAG signal at transcription start sites (left) and gene bodies (right) after treatment with either 50 nM dTAG-13, 2 μM revumenib, or 2 μM EZM-0414 for 48 hours.
